# Sensitive Fluorescent Biosensor Reveals Differential Subcellular Regulation of PKC

**DOI:** 10.1101/2024.03.29.587373

**Authors:** Qi Su, Jing Zhang, Wei Lin, Jin-Fan Zhang, Alexandra C. Newton, Sohum Mehta, Jing Yang, Jin Zhang

**Author notes:** Corresponding author: Zhang, Jin.

## Abstract

The protein kinase C (PKC) family of serine/threonine kinases, which consist of three distinctly regulated subfamilies, have long been established as critical for a variety of cellular functions. However, how PKC enzymes are regulated at different subcellular locations, particularly at emerging signaling hubs such as the ER, lysosome, and Par signaling complexes, is unclear. Here, we present a sensitive Excitation Ratiometric (ExRai) C Kinase Activity Reporter (ExRai-CKAR2) that enables the detection of minute changes in subcellular PKC activity. Using ExRai-CKAR2 in conjunction with an enhanced diacylglycerol (DAG) biosensor capable of detecting intracellular DAG dynamics, we uncover the differential regulation of PKC isoforms at distinct subcellular locations. We find that G-protein coupled receptor (GPCR) stimulation triggers sustained PKC activity at the ER and lysosomes, primarily mediated by Ca^2+^ sensitive conventional PKC (cPKC) and novel PKC (nPKC), respectively, with nPKC showing high basal activity due to elevated basal DAG levels on lysosome membranes. The high sensitivity of ExRai-CKAR2, targeted to either the cytosol or Par-complexes, further enabled us to detect previously inaccessible endogenous atypical PKC (aPKC) activity in 3D organoids. Taken together, ExRai-CKAR2 is a powerful tool for interrogating PKC regulation in response to physiological stimuli.

## 1. Introduction

Protein Kinase C (PKC) comprises a family of enzymes that play critical roles in regulating a variety of cellular processes, including cell growth, differentiation, and death. Aberrant PKC signaling drives many diseases, such as cancer, diabetes, and neurodegeneration^1–4^. Loss-of-function (LOF) mutations of PKC^1,2^, including LOF fusion oncoproteins^5^, frequently lead to cancer. Various PKC isoforms also play critical roles in synaptic plasticity, with isoform-selective knockout mice showing major deficits in brain functions^6,7^. On the other hand, even a minute elevation of PKCα activity by as little as 20% can impair cognitive function in an Alzheimer’s mouse model^4^. Thus, cells must be able to precisely balance the activity of PKC, as either too little or too much PKC activity can give rise to pathological states. Targeting specific PKC isoforms and their aberrant functions could promote new therapeutic efforts toward unmet needs^8^.

The nine PKC isoforms are categorized into three families: conventional PKC (cPKC), novel PKC (nPKC), and atypical PKC (aPKC). cPKC isoforms are activated by the binding of both Ca^2+^ and diacylglycerol (DAG), whereas nPKCs are activated by DAG alone, as their C2 domains lack residues that coordinate Ca^2+^ binding, while their C1 domains exhibit higher affinity towards DAG^9^. aPKCs show the greatest divergence in that they not only lack functional C1 domains, but their C2 domains are also substituted with PB1 domains that mediate interactions with specific scaffold proteins to induce kinase activation^10^. Inside the cell, PKC activity is tightly regulated at different subcellular locations through the interplay of second messengers, scaffold proteins, upstream kinases, and phosphatases. In the traditional model, activation of G_αq_-coupled GPCRs induces plasma membrane phospholipase C (PLC) activity, triggering production of DAG and inositol trisphosphate (IP_3_), the latter of which further evokes ER Ca^2+^ release. Binding of Ca^2+^ and/or DAG recruits both cPKCs and nPKCs to the plasma membrane, which is the strongest site of cPKC activation^11–14^. However, emerging evidence suggests that PKCs are active at other subcellular locations, such as the Golgi^11^ and nucleus^15^, and that PKCs can impact biological functions at other subcellular organelles. For instance, ER stress^16,17^ or the accumulation of lipids at the ER were both shown to activate PKC^18^, and PKC is also implicated in lysosome biogenesis^19,20^. Yet it remains unclear how PKCs are regulated at the ER or lysosome. aPKCs are also essential in regulating epithelial cell polarity and tumor epithelial-mesenchymal transition (EMT) ^21,22^. The Par3/Par6/aPKC complex promotes tight junction and apical domain formation by phosphorylating various targets to exclude them from the apical domain^22–24^. However, the activity dynamics and regulation of aPKC at Par complexes during polarity formation remain elusive, especially in a 3D culture, hindered by a lack of appropriate tools^10,22,25^.

Genetically encoded fluorescent biosensors are powerful tools to monitor kinase activities and small molecule dynamics, among various cellular events, in situ. Such biosensors have previously been used to study PKC activity in living cells, understand kinase structure-function relationships, and interrogate PKC regulation in various diseases^26,27^. The first Förster resonance energy transfer (FRET)-based C-kinase activity reporter (CKAR) was developed about two decades ago, comprising a PKC-specific substrate peptide and phosphoamino acid-binding FHA2 domain as the sensing unit sandwiched between a cyan/yellow fluorescent protein (FP) FRET pair^28^. CKAR has provided numerous insights into the regulation of PKC isoforms at different subcellular organelles. For example, Golgi-localized CKAR revealed sustained PKC activity by Ca^2+^-induced DAG synthesis at the Golgi^11^. Similarly, isoform-specific CKARs illuminated several long-overlooked aspects of PKC isoform regulations, such as Src kinase-controlled nuclear PKCδ activity^15^ and agonist-evoked aPKC activity^29,30^. However, FRET-based biosensors are limited by their small dynamic ranges^15,25^, and minute activities, such as that of endogenous aPKC in a 3D culture, have remained beyond the reach of these tools^10^.

Building on a newly developed excitation-ratiometric kinase activity reporter (ExRai-KAR) design^31–33^, we report a new excitation-ratiometric PKC activity reporter (ExRai-CKAR2) that enables highly sensitive tracking of PKC activity dynamics in living cells. We demonstrate the sensitivity and utility of ExRai-CKAR2 by elucidating the regulation of PKC isoforms at the ER and lysosome and by measuring the endogenous, basal aPKC activity in polarized MDCK organoids and lumenogenesis-related aPKC activity at the Par-complex in HEK293T organoids. Our results highlight the versatility of this robust biosensor in deciphering the spatiotemporal regulation of PKC in highly physiologically relevant settings.

## 2. Results

### Development and Characterization of ExRai-CKAR2

Our excitation-ratiometric kinase activity reporter (ExRai-KAR) series consist of circularly permutated EGFP (cpEGFP) sandwiched between a kinase-specific substrate peptide and phosphoamino acid-binding FHA1 domain. These biosensors exhibit two excitation peaks – one at ∼400 nm and another at ∼509 nm, with a distinct shoulder at ∼480 nm, corresponding to the neutral and anionic states of the GFP chromophore^34^ – along with a single emission peak at ∼515 nm^35^. Phosphorylation of the substrate peptide by the target kinase triggers binding by the FHA1 domain, with the resulting conformational change leading to increased fluorescence emission by the anionic species (e.g., 480-nm excitation) and decreased fluorescence emission by the neutral species (e.g., 400-nm excitation) (Figure 1a). The ratio of the fluorescence intensity induced by excitation at these two wavelengths (i.e., 480/400 excitation ratio) is thus used as a readout of kinase activity. Previously, we succeeded in generating an excitation-ratiometric PKC sensor, ExRai CKAR, by incorporating an established PKC-specific substrate peptide into this design^35^. To improve the performance of ExRai-CKAR, we took advantage of our recent success optimizing ExRai-KARs^31–33^ and mutated the two residues immediately proceeding and following cpEGFP in ExRai-CKAR. We tested several linker combinations and compared the responses of biosensor variants in HeLa cells stimulated with phorbol 12-myristate 13-acetate (PMA) to identify a sensor with an increased PKC-stimulated excitation ratio change (Supplementary Figure 1a). Of the biosensors tested, the variant with the linker pair SY-IS showed the largest response (Supplementary Figure 1b) and was designated ExRai-CKAR2 (Figure 1a).

**Figure 1.**
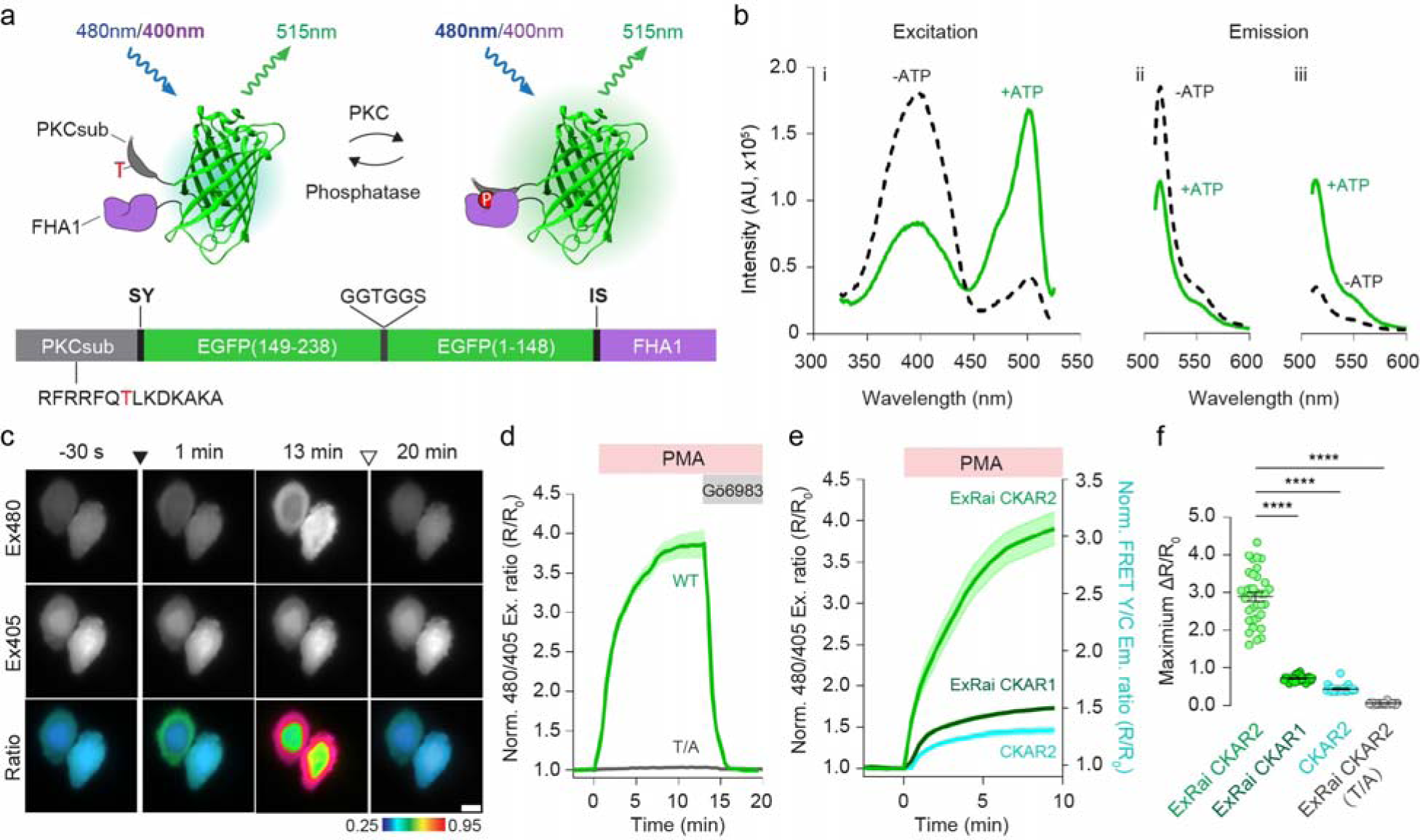
Development and characterization of ExRai-CKAR2. **a**, Modulation of cpEGFP fluorescence by a molecular switch dependent on PKC-mediated phosphorylation (top) and domain structure of ExRai-CKAR2 (bottom). **b**, Representative *in vitro* ExRai-CKAR2 fluorescence spectra collected at 530 nm emission (i) and 405 nm (ii) or 480 nm excitation (iii) without (black traces) or with (green traces) ATP, Ca^2+^, and lipids in the presence of purified PKCα. n = 3 independent experiments. **c**, Representative images of ExRai-CKAR2 fluorescence in Cos7 cells at 480-nm (Ex480, top) and 405-nm (Ex405, middle) excitation, along with pseudocolored images (bottom) of the excitation ratio change upon PMA stimulation and Gö6983 inhibition. Warmer colors indicate higher ratios. The solid arrowhead indicates PMA addition, and the hollow arrowhead indicates Gö6983 addition. Images are representative of three independent experiments. Scale bar, 10 μm. **d**, Representative average time courses showing the 480/405 excitation ratio responses of ExRai-CKAR2 (WT) and a nonphosphorylatable ExRai-CKAR2 T/A mutant (T/A) in Cos7 cells treated with PMA and Gö6983. **e**, Representative average time courses comparing the 480/405 excitation ratio or yellow/cyan emission ratio responses of ExRai-CKAR2, ExRai-CKAR1, or CKAR2 in Cos7 cells treated with 50 ng/ml PMA. **f**, Quantification of the maximum PMA-stimulated response of each biosensor. n = 35, n = 27, n = 20 and n = 22 cells from three independent experiments each. Time courses in **d** and **e** are representative of three independent experiments; solid lines indicate mean responses; shaded areas, s.e.m. Data in **f** were analyzed using ordinary one-way ANOVA followed by Dunnett’s multiple-comparison test. ****, *P* < 0.0001. Data are mean ± s.e.m.

We characterized the new ExRai-CKAR2 *in vitro* using purified protein. Purified ExRai-CKAR2 protein displayed two excitation maxima at approximately 400 nm and 500 nm, with a single emission peak at 516 nm (Figure 1b). *In vitro* phosphorylation by PKCα in the presence of ATP decreased the 405 nm peak by 38% and increased the 480 nm peak by 328%, compared to the no ATP/no Ca^2+^ control, resulting in a 433% increase in the 480 nm/405 nm excitation ratio (ΔR/R_0_).

In Cos7 cells, ExRai CKAR2 showed a 289 ± 12% increase in the 480/400 excitation ratio (ΔR/R_0_, mean ± s.e.m.; n = 35 cells) upon PMA stimulation (Figure 1c-e), representing a significant enhancement over both the second-generation FRET-based sensor CKAR2^36^ (R: FRET/CFP; ΔR/R = 46% ± 5%, n = 20 cells) and ExRai-CKAR1^35^ (72% ± 1%, n = 27 cells) (Figure 1e-f). The slightly reduced response amplitude compared to *in vitro* data might arise from different wavelength/filter choices of a fluorometer versus a microscope and phosphatase activity in cells^28^, but is consistent with a large dynamic range of the new biosensor. The addition of Gö6983, a potent pan-PKC inhibitor, reversed the PMA-induced responses to the baseline level (Figure 1c-d). In control experiments, Cos7 cells expressing a non-phosphorylatable mutant sensor (ExRai-CKAR2 T/A) showed no response to PMA and Gö6983 treatment (Figure 1c-d, Supplementary Figure 2). ExRai-CKAR2 also did not respond to activation of other AGC family kinases, as neither activation of PKA using Forskolin and 3-isobutyl-1-methylxanthine (IBMX) nor stimulation of Akt activity using PDGF in the presence of the PKC inhibitor Gö6983, evoked a detectable change in ExRai-CKAR2 signal (Supplementary Figure 3a-b)^28,31,32^. Additionally, stimulating Erk activity using epidermal growth factor (EGF) or AMPK activity using 2-deoxy glucose (2-DG) similarly failed to evoke detectable ExRai-CKAR2 responses (Supplementary Figure 3c-d), confirming that ExRai-CKAR2 specifically reports PKC activity. Compared to CKAR2 and ExRai-CKAR1, ExRai-CKAR2 is more sensitive, as shown by a broader PMA dose-response curve (Supplementary Figure 4). ExRai-CKAR2 responded to PMA doses as low as 0.1 ng/ml at endogenous PKC expression levels in Cos7 cells (ΔR/R = 2.8% ± 0.5%, n = 33 cells, *P* < 0.0001), whereas ExRai-CKAR1 only started to respond at doses above 0.5 ng/ml (ΔR/R_0_ = 1.6% ± 0.6%, n = 24 cells, *P* = 0.016) and CKAR2 responded at 5 ng/ml (ΔR/R_0_ = 12% ± 2%, n = 20 cells, *P* < 0.0001), suggesting a 5-to-50-fold sensitivity improvement with ExRai-CKAR2. The improved sensitivity and broad response curve allowed us to detect both subtle and strong PKC activities. Overall, ExRai-CKAR2 exhibited high selectivity as well as enhanced sensitivity and dynamic range.

### Sensitive Detection of Compartmentalized PKC Activity Using ExRai-CKAR2

PKC is present not only at the plasma membrane but also at various subcellular locations. Yet the regulation of different PKC isoforms at several emerging subcellular locations is not well understood. To enable sensitive detection of subcellular PKC activity, we targeted ExRai-CKAR2 to specific subcellular regions through the addition of established targeting motifs (Figure 2 and Supplementary Figure 5): a nuclear export signal (NES) for cytosolic targeting (Cyto-ExRai-CKAR2); a lipid modification motif derived from Lyn kinase (Lyn) for plasma membrane targeting (PM-ExRai-CKAR2); a cytochrome P450 (CYP450)-derived sequence for ER targeting (ER-ExRai-CKAR2); and full-length lysosome-associated membrane protein 1 (LAMP1)-derived sequence for lysosomal targeting (Lyso-ExRai-CKAR2).

**Figure 2.**
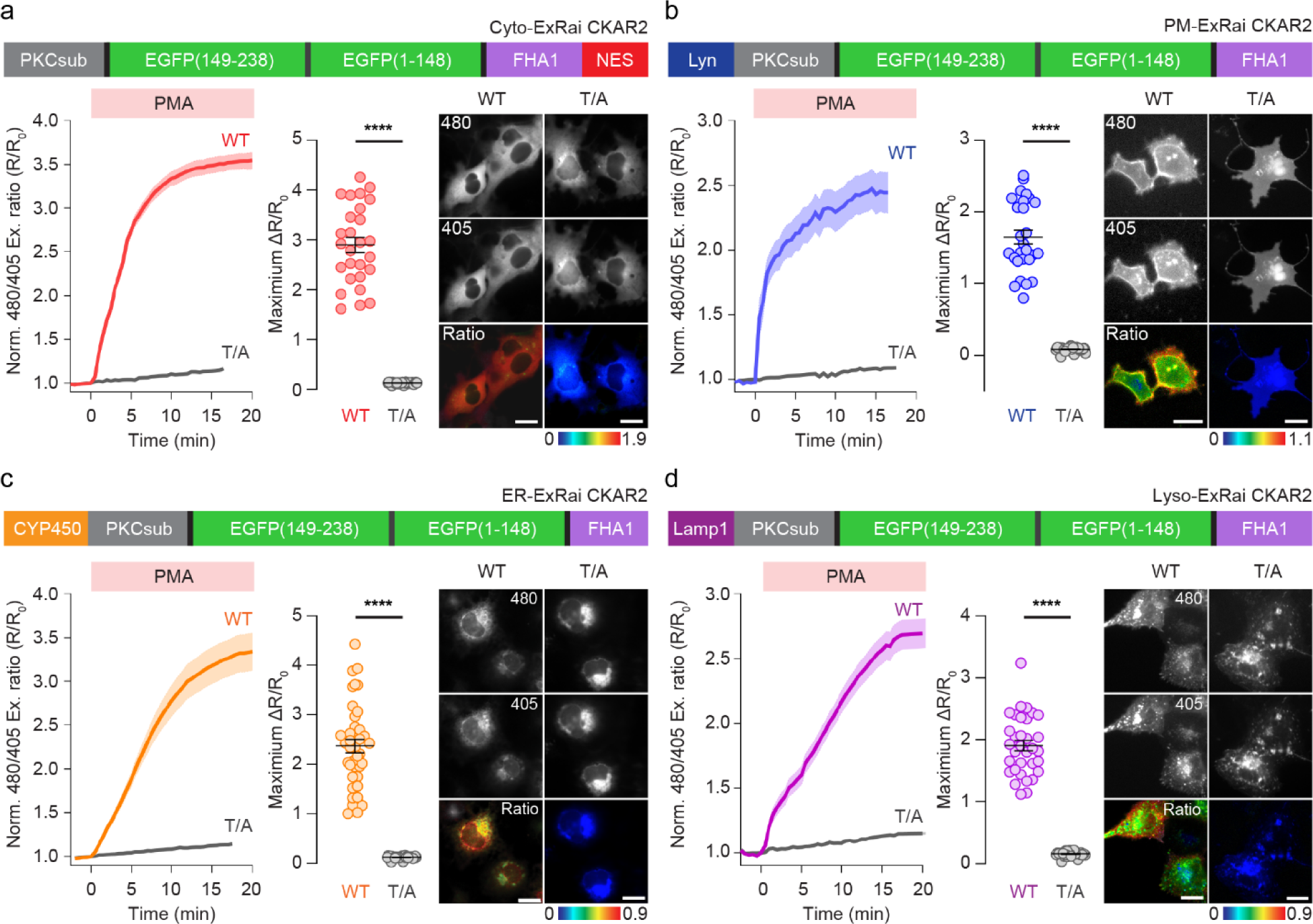
Subcellular targeting of ExRai-CKAR2. Domain structures (top), average excitation-ratio time courses of normalized excitation ratios (Ex480/405) (lower left), quantification of maximum response (middle right), and representative images (bottom right) of ExRai-CKAR2 and ExRai-CKAR2 T/A negative control targeted to **a**, cytoplasm (WT, n = 27 cells; T/A, n = 30 cells), **b**, plasma membrane (WT, n = 27 cells; T/A, n = 30 cells), **c**, ER (WT, n = 39 cells; T/A, n = 30 cells), and **d**, lysosomes (WT, n = 33 cells; T/A, n = 30 cells) in Cos7 cells stimulated with 50 ng/mL PMA. Scale bars, 10 μm. Time courses are representative of three independent experiments; solid lines indicate mean responses; shaded areas, s.e.m. Statistical analyses ere performed using Student’s *t*-test. \*\*\*\**P* < 0.0001. Data are mean ± s.e.m.

Upon PMA stimulation, Cyto-ExRai-CKAR2 and PM-ExRai-CKAR2 reported 290% ± 15% (ΔR/R_0_, n = 30 cells, Figure 2a) and 144% ± 16% (ΔR/R_0_, n = 24 cells, Figure 2b) increases in the 480 nm/400 nm excitation ratio in the cytosol and at the plasma membrane, respectively, consistent with well-characterized PKC activity at these locations. Conversely, no response was detected using ExRai-CKAR2 T/A targeted to either location (Figure 2a, b). ER-ExRai-CKAR2 and Lyso-ExRai-CKAR2 also revealed PMA-induced PKC activity at the ER and lysosome membranes, two previously under-appreciated sites of PKC signaling. ER-ExRai-CKAR2 responded to PMA stimulation with a 234% ± 22% increase in the 480 nm/405 nm excitation ratio (ΔR/R_0_; n = 26 cells), whereas the T/A mutant showed no PMA-induced response (Figure 2c). Similarly, Lyso-ExRai-CKAR2 exhibited a 178% ± 12% excitation ratio change (ΔR/R_0_, n = 30 cells) upon PMA stimulation, whereas no response was detected using the T/A mutant sensor (Figure 2d). In summary, ExRai-CKAR2 enables sensitive and robust detection of PKC activity at various subcellular locations.

### PKC regulation at the ER and lysosome

Although traditionally regarded as organelles dedicated to overseeing protein trafficking, quality control, and degradation, the ER and lysosome have emerged as key intracellular signaling hubs^37–39^. The detection of ER and lysosomal PKC activity using targeted ExRai-CKAR2 is consistent with past reports that PKCα accumulates on the ER surface upon phorbol ester activation in 3T3 cells^40^ and that PKC contributes to lysosome biogenesis, with PMA stimulation inducing PKC localization to lysosomes^19^. However, the regulation of PKC activity at these sites in response to physiological signals has not been explored. We therefore set out to investigate ER and lysosomal PKC regulation in response to UTP stimulation of the G_αq_-coupled P2Y receptor.

As shown in Figure 3a, stimulating Cos7 cells expressing ER-ExRai-CKAR2 or Lyso-ExRai-CKAR2 with UTP (100 μM) induced clear elevations in PKC activity at each location, with a 21% ± 2% excitation ratio increase (ΔR/R_0_, n = 36 cells) from ER-ExRai-CKAR2 and a 40% ± 2% excitation ratio increase (ΔR/R_0_, n = 36 cells) from Lyso-ExRai-CKAR2. We next assessed the dynamics of agonist-induced PKC activity at these two locations by quantifying the fraction of sensor response remaining at 15 min after stimulation versus the maximum response (i.e., sustained activity metric at 15 min, or SAM15)^41,42^. Interestingly, both ER and lysosomal PKC activities were sustained after UTP stimulation, with a SAM15 of 0.65 ± 0.02 (n = 36 cells) at the ER and a SAM15 of 0.92 ± 0.05 (n = 36 cells) at the lysosome (Figure 3a). By contrast, UTP stimulation induced a sharp, 75% ± 3% (ΔR/R_0_, n = 25 cells) excitation ratio increase from PM-ExRai CKAR2, followed by fast reversal (SAM15 = 0.06 ± 0.05, n = 25 cells; *P* < 0.0001 vs. ER or lysosome) (Figure 3a), consistent with previous findings that G_αq_ signaling mediates fast and transient PKC activity at the plasma membrane^11^. Notably, treatment with the pan-PKC inhibitor Gö6983 following UTP stimulation led to an incomplete reversal of the ER-ExRai-CKAR2 response compared with Lyso-ExRai-CKAR2 (Supplementary Figure 6), suggesting a lower level of phosphatase activity at the ER.

**Figure 3.**
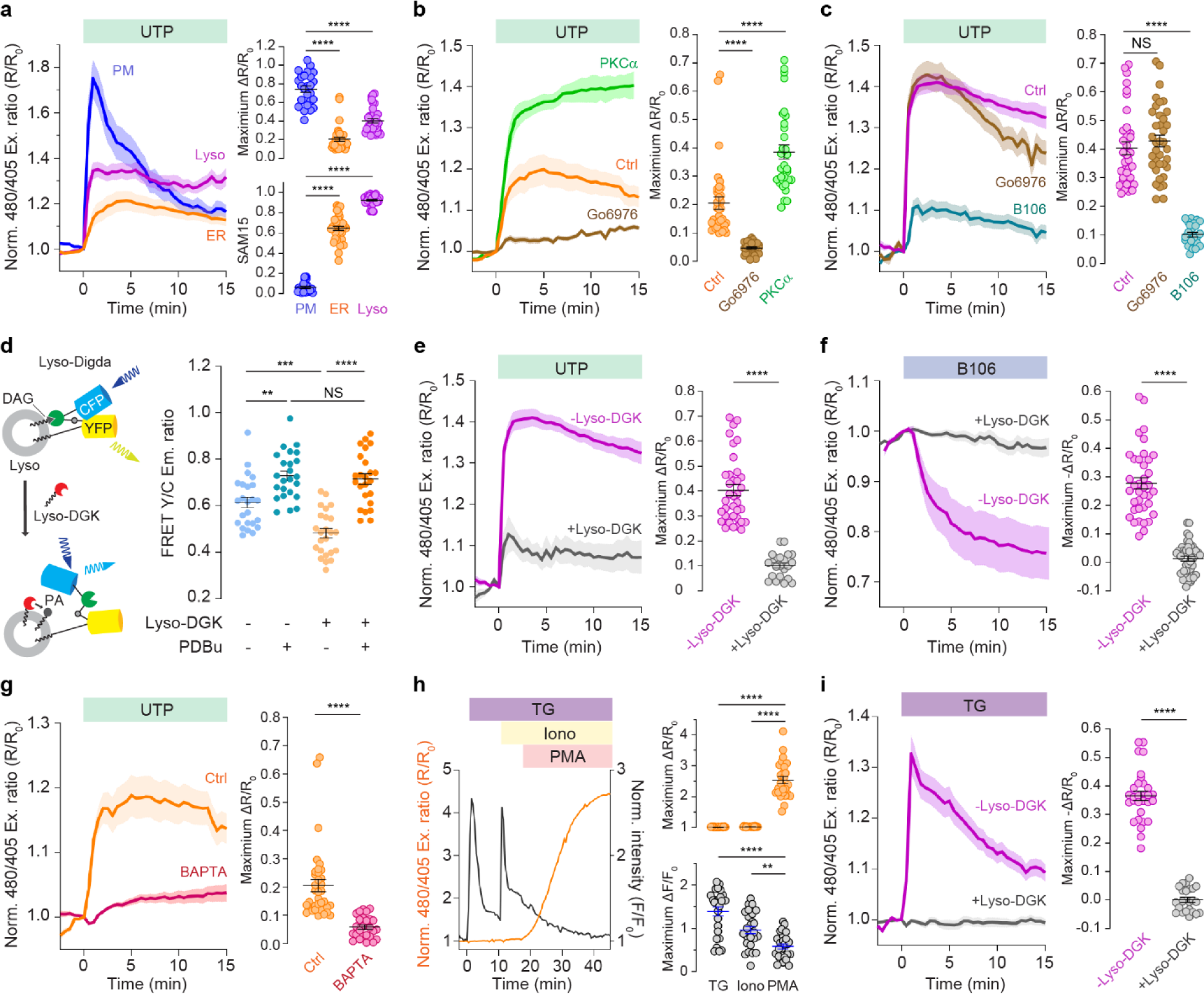
PKC regulation at the ER and lysosome. **a**, Representative average time courses (left) of normalized excitation ratios (Ex480/405) from PM-ExRai-CKAR2 (PM, n = 10 cells), ER-ExRai-CKAR2 (ER, n = 15 cells), and Lyso-ExRai-CKAR2 (Lyso, n= 16 cells) in Cos7 cells expressing stimulated with 100 μΜ UTP. Maximum responses (ΔR/R_0_, top right) and SAM15 (bottom right) are quantified from n = 25, n = 36 and n = 36 cells from three independent experiments. **b**, Representative average time courses (left) of normalized excitation ratio (Ex480/405) responses to 100 μΜ UTP stimulation in Cos7 cells expressing ER-ExRai-CKAR2 without (Ctrl, n = 10 cells) of with (Go6976, n = 12 cells) 30 min pre-incubation with the cPKC inhibitor Go6976, or co-overexpressed with PKCα-mCherry (PKCα, n= 14 cells). Maximum responses (right) quantified from n = 36, n = 34 and n = 32 cells from three independent experiments. **c**, Representative average time courses (left) of normalized excitation ratio (Ex480/405) responses 100 μΜ UTP to treatment in Cos7 cells expressing Lyso-ExRai CKAR2 without (Ctrl, n = 12 cells) or with 30-min pretreatment (Go6976) with the cPKC inhibitor Go6976 (Go6976, n = 16 cells) or thenPKC inhibitor (5 μM) B106 (B106, n = 9 cells). Maximum responses (right) quantified from n = 36, n = 37 and n = 24 cells from three independent experiments. **d**, Schematic illustrating depletion of lysosomal DAG levels using Lyso-DGK (left). Raw yellow/cyan emission ratios from Lyso-Digda-expressing Cos7 cells with or without PDBu stimulation or Lyso-DGK co-expression (right). n = 24, n = 24, n = 24 and n = 24 cells from three independent experiments. *P* = 0.0015 (-lyso-DGK/-PDBu vs -lyso-DGK/+PDBu), *P* = 0.0002 (-lyso-DGK/-PDBu vs +lyso-DGK/-PDBu); ordinary one-way ANOVA followed by Tukey’s multiple-comparison test. **e**, Representative average time courses (left) of normalized excitation ratio (Ex480/405) responses stimulation with 100 μΜ UTP in Cos7 cells expressing Lyso-ExRai-CKAR2 without (-lyso-DGK, n = 17 cells) or with (+lyso-DGK, n = 10 cells) Lyso-DGK co-expression. Maximum responses (right) quantified from n = 36 and n = 22 cells from three independent experiments. **f**, Representative average time courses (left) of normalized excitation ratios (Ex480/405) in Cos7 cells expressing lyso-ExRai CKAR2 (-lyso-DGK, n = 18 cells) or lyso-ExRai CKAR2 plus lysosomal DAG kinaseζ (+lyso-DGK, n = 20 cells) treated with 5 μΜ nPKC inhibitor B106. Quantification of maximum response (right). n = 38 and n = 44 cells from three independent experiments. **g**, Representative average time courses (left) of normalized excitation ratio (Ex480/405) responses to stimulation with 100 μΜ UTP in ER-ExRai-CKAR2-expressing Cos7 cells without (Ctrl, n = 15 cells) or with (BAPTA, n = 11 cells) 30-min pretreatment with the Ca^2+^ chelator BAPTA-AM (20 μM). Maximum responses (right) quantified from n = 36 and n = 26 cells from three independent experiments. **h**, Representative single-cell time courses of normalized excitation ratio (Ex480/405) (orange curve) or normalized fluorescent intensity (black curve) response in Cos7 cells co-expressing ER-ExRai-CKAR2 and RCaMP and sequentially stimulated with 1 μM thapsigargin (TG), 1 μM ionomycin (Iono), and 50 ng/ml PMA. Maximum response to each treatment (right) quantified from n = 26 cells from three independent experiments. **i**, Representative average time courses (left) of normalized excitation ratios (Ex480/405) in Cos7 cells expressing lyso-ExRai CKAR2 (-lyso-DGK, n = 10 cells), or lyso-ExRai CKAR2 co-overexpressed with lyso-DGK (+lyso-DGK, n = 9 cells) stimulated by thapsigargin (TG). Quantification of maximum response (right). n = 30 and n = 23 cells from three independent experiments. All time courses are representative of three independent experiments; solid lines indicate mean responses; shaded areas, s.e.m. For **a-c** and **i**, statistical analysis was performed using ordinary one-way ANOVA followed by Dunnett’s multiple-comparison test. For **e**, **f, g** and **i**, statistical analysis was performed using Student’s *t*-test. \*\**P* < 0.01, \*\*\**P* < 0.001, \*\*\*\**P* < 0.0001. Data are mean ± s.e.m.

We then investigated the contribution of different PKC isoforms to the PKC activity that we measured on the ER and lysosome surfaces. Pretreating Cos7 cells with 1 μM Go6976, a cPKC-specific inhibitor, prior to stimulation with UTP, greatly reduced the response from ER-ExRai-CKAR2 (ΔR/R_0_: 4.7% ± 0.4%; n = 34 cells, *P* < 0.0001) (Figure 3b), suggesting that cPKC accounts for roughly 3/4 of the receptor-mediated PKC response at the ER. Consistent with this result, overexpressing PKCα, a typical cPKC isoform, strongly enhanced the UTP-stimulated response from ER-ExRai-CKAR2 to 39% ± 3% (n = 32 cells, *P* < 0.0001, Figure 3b). On the other hand, Go6976 pretreatment had no effect on the maximum UTP-stimulated response of Lyso-ExRai CKAR2 (ΔR/R_0_: 43% ± 2%, n = 37 cells, *P* = 0.403) compared to control cells (Figure 3c). Instead, we found that pretreatment with 5 μM B106, an nPKC-specific inhibitor^43^, significantly decreased the UTP-induced lysosomal PKC activity reported by Lyso-ExRai-CKAR2 (ΔR/R_0_: 10.1% ± 0.8%; n = 24 cells, *P* < 0.0001, Figure 3c), suggesting a more critical role for nPKC in controlling PKC activity at the lysosome.

Surprisingly, we also found that B106 addition induced a 28% ± 2% (ΔR/R_0_, n = 38 cells) decrease in the Lyso-ExRai-CKAR2 excitation ratio on its own (Figure 3f), suggesting the presence of high basal nPKC activity on the lysosome surface. Second messengers play vital roles in regulating PKC isoforms, and given that nPKCs are solely dependent on DAG for activation, we hypothesized that this basal nPKC activity is supported by basal accumulation of DAG at lysosomes. However, little is known about lysosomal DAG dynamics or whether DAG is even present in lysosome membranes. We therefore generated a lysosome-targeted DAG biosensor to help test our hypothesis. Starting with the existing, PM-targeted DAG biosensor Digda^44^, we replaced the C-terminal PM targeting motif with full-length LAMP1 for lysosomal targeting, and employed an enhanced cyan fluorescent protein Cerulean as the FRET donor (Supplementary Figure 7a). The resulting sensor, designated Lyso-Digda, colocalized with LysoTracker Red-positive puncta in Cos7 cells (Pearson’s coefficient R = 0.77 ± 0.05, n = 17, Supplementary Figure 7b) and showed a 21% ± 1% yellow/cyan emission ratio increase (ΔR/R_0_, n = 35 cells) upon treatment with the phorbol ester Phorbol 12,13-dibutyrate (PDBu) (Supplementary Figure 7c). Consistent with our Lyso-ExRai-CKAR2 results, cells stimulated with 100 μM UTP also showed a 9.4% ± 0.6% (ΔR/R_0_, n = 39 cells) emission ratio increase from Lyso-Digda (Supplementary figure 7c).

To examine whether DAG basally accumulates at lysosomes, we then applied a targeted biochemical perturbation approach to selectively alter DAG levels on the lysosome membrane^32,45^. Diacylglycerol kinase ζ (DGKζ) catalyzes the conversion of DAG to phosphatidic acid (PA) and should be able to deplete basal DAG levels and suppress receptor-stimulated DAG accumulation at target sites (Figure 3d). Indeed, when we targeted mRuby2-fused DGKζ to the lysosome using the LAMP1 motif (Lyso-DGK), we recorded a significant reduction in the basal emission ratios from Lyso-Digda (R = 0.48 ± 0.10) in cells co-expressing Lyso-DGK versus control cells (R = 0.61 ± 0.11; *P* < 0.0001, Figure 3d). However, treatment with the non-degradable phorbol ester PDBu increased the Lyso-Digda emission ratio to the same level in both the presence (R = 0.73 ± 0.10) and absence (R = 0.72 ± 0.11; *P* = 0.633) of Lyso-DGK (Figure 3d), suggesting that Lyso-Digda can measure both basal and stimulated DAG levels at the lysosome and that Lyso-DGK effectively depleted high basal DAG at lysosomes.

Looking at PKC activity, we found that Lyso-DGK co-expression suppressed both the UTP-stimulated increase (ΔR/R_0_ = 10% ± 1%, n = 22 cells, *P* < 0.0001, Figure 3e) and the B106-induced decrease (ΔR/R_0_ = 1.3% ± 0.8%, n = 44 cells, *P* < 0.0001 vs no Lyso-DGKζ, Figure 3f) in the Lyso-ExRai-CKAR2 excitation ratio. These data support our hypothesis that a basal pool of lysosomal DAG drives elevated nPKC activity on the lysosome surface in the absence of GPCR signaling.

Unlike nPKCs, cPKC activation requires both DAG and Ca^2+^-binding. We therefore hypothesized that Ca^2+^ is essential for the induction of ER PKC activity. Indeed, when we pretreated Cos7 cells with 20 μM BAPTA-AM to chelate intracellular Ca^2+^, we observed a dramatic reduction in the UTP-stimulated ER-ExRai-CKAR2 response (ΔR/R: 5.8% ± 0.8%; n = 26 cells, *P* < 0.0001, Figure 3g). Interestingly, treatment with thapsigargin (TG, 1 μM), a potent blocker of ER Ca^2+^-ATPases, failed to elicit a response from ER-ExRai-CKAR2 (ΔR/R: 1% ± 1%, n = 26 cells), despite clearly evoking a robust response from the red-fluorescent Ca^2+^ indicator RCaMP^46^ (Figure 3h). Even global extracellular Ca^2+^ influx triggered by the addition of 1 μM ionomycin was not capable of inducing a response from ER-ExRai-CKAR2 (ΔR/R: 3.9% ± 0.9%, n = 26 cells; Figure 3h), suggesting that Ca^2+^ is necessary, but not sufficient, for PKC activity at the ER. Conversely, TG induced a robust 37% ± 2% increase in the Lyso-ExRai-CKAR2 excitation ratio (ΔR/R_0_, n = 30 cells, Figure 3i), suggesting that elevated Ca^2+^ is sufficient to trigger lysosomal PKC activity. This effect did not require the presence of increased DAG levels at the lysosome, as Lyso-Digda showed no response to TG stimulation (ΔR/R_0_ = 0.6% ± 0.5%, n = 21 cells, Supplementary Figure 7c). However, basal DAG enrichment on the lysosome membrane appeared to be critical for this Ca^2+^-induced lysosomal PKC activity, as overexpressing Lyso-DGK completely suppressed the TG-stimulated Lyso-ExRai-CKAR2 response (ΔR/R_0_ = 0.1% ± 0.9%, n = 23 cells, *P* < 0.0001) (Figure 3i). Given that nPKCs are not directly responsive to Ca^2+^, our data suggests that cPKC can also play a critical role in mediating PKC activity at the lysosome. Indeed, when we pretreated cells with the cPKC-selective inhibitor Go6976, the TG-induced lyso-ExRai CKAR2 response was significantly reduced (ΔR/R_0_ = 11% ± 1%, n = 24 cells, Supplementary Figure 8) compared with no pretreatment. Taken together, these data show that PKC exhibits complex subcellular regulation that is dependent on subcellular localization and stoichiometry of isoforms, basal level and spatiotemporal dynamics of second messengers.

### aPKC regulation in polarized or lumenized 3D organoid models

aPKCs are unique among PKC isoforms in that they are regulated by neither DAG nor Ca^2+^, but rather activated through binding to scaffold proteins. Binding of aPKC with a scaffold protein relieves autoinhibition by pulling the pseudosubstrate away from the substrate binding cavity^10,47^. For example, a major aPKC isoform, PKCζ, is activated and kept in an open, active conformation through interactions with the cell polarity-associated protein Par6. The PKCζ-containing Par complex is an essential regulator of apical-basal polarity in epithelial cells^22^. aPKC regulates apical domain formation by phosphorylating and controlling various targets to specific cell domains, such as apical vs. basal domains^23,24^. Loss of apical-basal polarity has been shown to downregulate PKCζ-mediated phosphorylation of SNAI1 to promote epithelial-mesenchymal transition (EMT)^22^. Direct interrogation of aPKC activity during epithelial polarization should thus provide critical insights into molecular and cellular processes involved in cancer metastasis. However, the modest activity of aPKC complicates efforts to directly measure endogenous aPKC activity in living cells.

Therefore, we tested the capability of ExRai-CKAR2 to detect aPKC activity by co-overexpressing an N-terminus truncated, constitutively active aPKC isoform, PKMζ, with ExRai CKAR2 or the phosphorylate site mutant (T/A) in HeLa cells. To visualize basal aPKC activity, we directly treated HeLa cells expressing different versions of CKAR with the aPKC-selective inhibitor pz09^48^ and quantified the subsequent decrease in biosensor signal. With pz09 treatment, ExRai-CKAR2 showed a 28.2% ± 0.9% decrease in excitation ratio (ΔR/R_0_, n = 24 cells, *P* < 0.0001 compared to CKAR2), whereas the negative control ExRai-CKAR2 T/A showed no response (ΔR/R_0_: 0.1% ± 0.1%, n = 28 cells) (Figure 4b). Strikingly, in the absence of PKMζ overexpression, ExRai-CKAR2-showed a 12.4% ± 0.9% (ΔR/R_0_, n = 24 cells) excitation ratio decrease in HeLa cells treated with 5 μM pz09, demonstrating the capability of the reporter to detect endogenous aPKC activity (Figure 4a&c), whereas CKAR2, the previous best-performing FRET-based PKC sensor, failed to capture the endogenous basal aPKC activity (ΔR/R_0_ = 0.8% ± 0.4%, n = 20 cells, *P* = 0.165) (Figure 4a and c).

**Figure 4.**
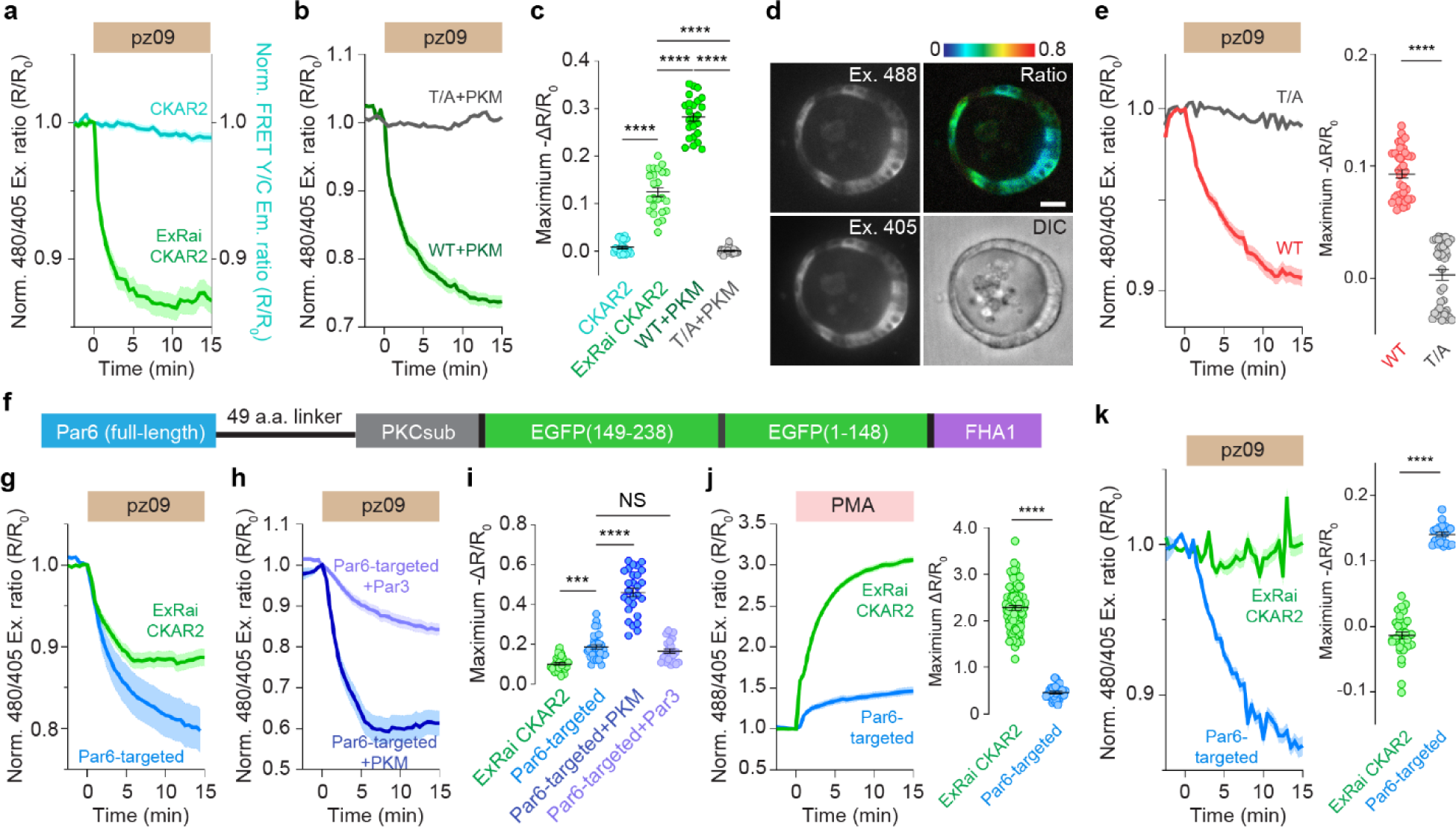
ExRai-CKAR2 reports endogenous aPKC activity in organoids. **a**, Representative average time courses of normalized excitation ratio (Ex480/405) or yellow/cyan emission ratio responses in HeLa cells expressing ExRai-CKAR2 (n = 24 cells) or CKAR2 (n = 20 cells), respectively, treated with 5 μM pz09. **b**, Representative average time courses of normalized excitation ratio (Ex480/405) responses to 5 μM pz09 addition in HeLa cells co-overexpressing mCherry-tagged constitutively active PKMζ along with ExRai CKAR2 (WT+PKM, n = 24 cells) or the negative control ExRai-CKAR2 T/A (T/A+PKM, n = 28 cells). **c**, Maximum responses from **a** and **b** quantified from n = 20, n = 24, n = 24 and n = 28 cells from three independent experiments. **d**, Representative fluorescence, pseudocolor, and DIC images of 3D-cultured MDCK organoids stably expressing cyto-ExRai-CKAR2. The pseudocolor image depicts raw Cyto-ExRai-CKAR2 excitation ratios (Ex480/405); warmer colors indicate higher ratios. Scale bar, 20 μm. **e**, Representative average time courses (left) of normalized excitation ratio (Ex480/405) responses to pz09 treatment in 3D-cultured MDCK organoids expressing Cyto-ExRai CKAR2 (WT, n = 38 cells) or Cyto-ExRai CKAR2 T/A (T/A, n = 40 cells). Maximum responses (right) quantified from n = 38 and n = 40 cells in 4 organoids. **f**, Domain structure of Par6-ExRai-CKAR2. **g**, Representative average time courses of normalized excitation ratios (Ex480/405) in HEK293T cells expressing ExRai-CKAR2 (n = 35 cells) or Par6-ExRai CKAR2 (n = 30 cells) treated with pz09. **h**, Representative average time courses of normalized excitation ratios (Ex480/405) responses to pz09 addition in HEK293T cells co-expressing Par6-ExRai-CKAR2 along with PKMζ-mCherry (+PKM, n = 28 cells) or Par3-mCherry (+Par3, n = 30 cells). **i**, Maximum response from **g** and **h** quantified from n = 28, n = 30, n = 28 and n = 23 cells from three independent experiments. *P* = 0.0005 (ExRai-CKAR2 versus Par6). **j**, Representative average time courses (left) of normalized excitation ratio (Ex480/405) response to PMA stimulation in HEK293T cells expressing ExRai-CKAR2 (n = 30 cells) or Par6-ExRai-CKAR2 (n = 20 cells). Maximum response (right) quantified from n = 80 and n = 32 cells from three independent experiments. **k**, Representative average time courses (left) of normalized excitation ratio (Ex480/405) response to pz09 treatment in 3D-cultured HEK293T organoids expressing ExRai-CKAR2 (n = 31 cells) or Par6-ExRai-CKAR2 (n = 20 cells). Maximum responses (right) quantified from n = 31 and n = 20 cells in 2 organoids. All time courses are representative of three independent experiments; solid lines indicate mean responses; shaded areas, s.e.m. Statistical analyses were performed using ordinary one-way ANOVA followed by Tukey’s multiple-comparison test (**c**, **i**) or Student’s *t*-test (**e**, **j**, **k**). \*\*\**P* < 0.001, \*\*\*\**P* < 0.0001. Data are mean ± s.e.m.

The ability of ExRai-CKAR2 to detect endogenous aPKC activity opens the door to studying aPKC regulation under more physiological conditions, such as during epithelial polarity formation. To explore this possibility, we first tested Cyto-ExRai-CKAR2 during MDCK cell polarization. Widely used to study epithelial polarity, MDCK cells can form polarized epithelial organoids with intact apical-basal polarity in 3D Matrigel culture. To validate our approach, we first treated 2D-cultured MDCK cells expressing Cyto-ExRai-CKAR2 with 5 μM pz09 and observed a 16% ± 1% excitation ratio decrease (ΔR/R_0_, n = 24 cells, Supplementary Figure 9), whereas cells expressing the negative control T/A mutant sensor showed no detectable response (ΔR/R_0_ = 0.7% ± 0.3%, n = 23 cells, Supplementary Figure 9). Based on these results, we next generated MDCK cells stably expressing Cyto-ExRai CKAR2, which we cultured in 3D. After 9 days of growth on Matrigel, MDCK cells developed into polarized organoids with the signature hollow lumen structure (Figure 4d). Excitation-ratiometric imaging of polarized MDCK organoids via spinning disc confocal microscopy revealed a robust, 9.3% ± 0.4% (ΔR/R_0_, n = 38 cells in 4 organoids) decrease in the Cyto-ExRai-CKAR2 excitation ratio upon addition of pz09. Conversely, MDCK organoids generated from cells stably expressing the negative control Cyto-ExRai-CKAR2 T/A showed no response to pz09 (ΔR/R_0_ = −0.3% ± 0.5%, n = 40 cells in 4 organoids, Figure 4e). Thus, ExRai-CKAR2 enables direct, real-time visualization of polarity-related, endogenous aPKC activity in a 3D culture model while maintaining single-cell resolution.

Although Cyto-ExRai-CKAR2 successfully reported aPKC activity upon pz09 treatment, we sought to further enhance the selectivity of aPKC activity detection by targeting ExRai-CKAR2 directly to the Par complex. Previous efforts using the PB1 domain of Par6 to target isoform-specific aCKAR were unsuccessful in localizing the biosensor to the Par complex, as images of cells expressing the fused biosensor lacked the signature punctate structure of the Par complex^10,49,50^. We therefore tested directly tethering ExRai-CKAR2 to full-length Par6 (Figure 4f). Our initial testing revealed that incorporating short linkers of only 2 or 8 residues between Par6 and ExRai-CKAR2 rendered the sensor poorly responsive to pz09 treatment (Supplementary Figure 10). We ultimately found that inserting a flexible, 49-aa linker between Par6 and ExRai-CKAR2 yielded a sensor with a robust response to pz09 addition. The resulting Par6-ExRai-CKAR2 sensor localized to punctate structures^50^ when expressed in HeLa cells and colocalized with other Par-complex components, namely, PKCζ (Pearson’s coefficient R = 0.93 ± 0.05, n = 12 cells) and Par3 (Pearson’s coefficient R = 0.82 ± 0.07, n = 12 cells) (Supplementary Figure 11). Par6-ExRai-CKAR2 showed a larger response (ΔR/R_0_ = −19% ± 1%, n = 30 cells, *P* = 0.0002, Figure 4g & i) than untargeted ExRai-CKAR2 upon pz09 treatment, demonstrating enhanced sensitivity for detecting endogenous aPKC activity. Upon overexpression of constitutively active PKMζ, pz09 induced an even stronger 46% ± 2% excitation ratio decrease (n = 27 cells, *P* < 0.0001, Figure 4h-i), whereas co-overexpression of Par3 did not affect the sensor response (ΔR/R_0_: −17% ± 1%, n = 23 cells, *P* = 0.229, Figure 4h-i). To rule out potential activation of aPKC by the targeting motif, Par6, we compared this result with a variant containing an A30D mutation in Par6, which was shown to disrupt aPKC binding and activation^10^. Par6(A30D)-ExRai-CKAR2 showed comparable response to pz09 (ΔR/R_0_: −16% ± 2%, n = 18 cells, *P* = 0.101, Supplementary Figure 12), suggesting Par6-ExRai-CKAR2 does not artificially activate aPKC. Strikingly, Par6-ExRai-CKAR2 also showed a significantly smaller response to PMA stimulation (ΔR/R_0_ = 47% ± 5%, n = 20 cells), which should lead to full activation of cPKC and nPKC, compared with untagged ExRai-CKAR2 (ΔR/R_0_ = 224%, *P* < 0.0001, Figure 4j). To quantify the selectivity of Par6-ExRai-CKAR3 towards aPKC over other PKC isoforms, we calculated the ratio of the pz09-induced (i.e., aPKC) response versus the PMA-induced (i.e., pan-PKC) response. Compared to Cyto-ExRai-CKAR2 (ΔR_pz09_/ΔR_PMA_ = 0.043), Par6-ExRai-CKAR2 (ΔR_pz09_/Δ R_PMA_ = 0.44) showed a marked, 10-fold greater selectivity towards aPKC, indicating a reduced potential for interference from other PKC isoforms in studies of native aPKC regulation.

Par6-ExRai-CKAR2 serves as an ideal tool to visualize aPKC regulation at the Par-complex. Melanoma cell adhesion molecule (MCAM), a membrane receptor endogenously expressed in HEK293 cells, has been shown to mediate apical-basal polarity-driven lumenogenesis in HEK293 organoids^51^. Although Cdc42, an upstream regulator of Par6-aPKC in polarity formation^52^, was shown to be polarized in this lumenogenesis model, aPKC activity has not been characterized. Using Par6-ExRai-CKAR2, we found that aPKC is active during HEK293T lumenogenesis (Supplementary Figure 13). In 3D-cultured HEK293T cells transiently expressing Par6-ExRai-CKAR2, pz09 addition induced a 14.1% ± 0.3% excitation ratio decrease (n = 20 cells in 2 organoids, Figure 4k). On the other hand, cells expressing untargeted ExRai-CKAR2 showed minimal responses upon pz09 treatment under the same culture conditions (ΔR/R_0_: 1.4% ± 0.6%, n = 31 cells in 2 organoids, Figure 4k). Together, these data demonstrate ExRai CKAR2, particularly, the Par-targeted variant, serves as a sensitive biosensor of aPKC activity.

## Discussion

Genetically encoded fluorescent kinase activity reporters are being increasingly relied on to study the spatiotemporal regulation of signaling pathways in various model systems, calling for a more extensive expansion of their versatility and advancement of their performance. Here, we introduced ExRai-CKAR2, the most sensitive PKC activity reporter thus far, with a roughly 57-fold higher dynamic range compared to CKAR1^28^, 6-fold higher dynamic range compared to CKAR2^36^ and 4-fold higher dynamic range compared to ExRai-CKAR1^35^. ExRai-CKAR2 enables us to detect subtle PKC activity changes at different subcellular locations and to monitor endogenous aPKC activity in organoid models, which posed a challenge to previous PKC activity biosensors. By tuning the spacer length between the targeting domain Par6 and ExRai-CKAR2, we were able to obtain Par6 targeted biosensors that retain similar dynamic ranges as the untargeted sensor (Figure 4i & Supplementary Figure 10), empowering our efforts to dissect spatiotemporal PKC regulation. Future engineering will focus on generating isoform-specific biosensors and further increasing dynamic range. Overall, ExRai-CKAR2 represents notable progress in detecting subcellular PKC activity with high sensitivity and facilitates robust visualization of subtle PKC signaling dynamics with subcellular precision, even in 3D cultures.

PKC isoforms exhibit different, sometimes opposing, functions under various physiological and pathological conditions, including cancer development^53^. This phenomenon has previously been attributed to cancer-or cell-type-specific differences^1,5,43,54^. Using subcellular targeted ExRai-CKAR2, we discovered that different PKC isoforms exhibit distinct subcellular activity patterns. Although we detected PKC activity at both the ER and lysosomes upon GPCR activation by UTP, we found that cPKCs were the primary contributor to ER-localized PKC activity, whereas nPKCs were the dominant isoform responsible for lysosomal PKC activity. Thus, PKC isoforms may drive specific functional effects by forming spatially distinct signaling territories.

The ability to sensitively detect subcellular PKC activities also enabled us to more closely investigate their regulation. The dominant role of cPKC in G_q_-stimulated PKC activity at the ER is consistent with the activation of cPKC by Ca^2+^ and DAG, as well as previous work showing that PKCα localizes to the ER. Imaging and mass spectrometry analyses^55,56^ also indicate that DAG is enriched on ER membranes in oocytes and muscle cells. On the other hand, a principal role for nPKC in lysosomal PKC activity is consistent with previous results showing the lysosomal localization of PKCδ^57^. Although phospholipase C (PLC), which catalyzes DAG production from phosphatidylinositol 4,5-bisphosphate (PI[4,5]P_2_), has been shown to localized to lysosomes and regulate lysosome stability^58,59^, few studies have directly investigated lysosomal DAG accumulation. We discovered that DAG is not only present but is also basally enriched on lysosome membranes, supporting both basal nPKC activity and Ca^2+^-induced cPKC activation at this location, partially mirroring Golgi PKC signaling^15,60^. However, how G_q_-mediated signaling induces lysosomal DAG elevations remains unclear. In contrast to the Golgi, however, Ca^2+^ did not directly induce lysosomal DAG synthesis, suggesting that G_q_-mediated DAG increases at the lysosome require additional regulators. We recently showed that the trafficking of active GPCR and Gα_s_ to endosomes is critical for GPCR-mediated Erk activation^61^. While G_q_-coupled GPCRs are also recycled through the endosome-lysosome pathway^62^, it remains to be seen whether they remain active at these locations to directly regulate lysosomal PLC.

Given their ability to mimic complex organs, cultured organoids are increasingly recognized as valuable *in vitro* models with huge clinical potential^63^. The importance of Par-aPKC complexes in cell polarity formation and EMT has been well documented in these models, with more than half of all known polarity-regulating proteins identified as aPKC substrates^22,24^. However, how aPKC is regulated by and functionally integrates upstream signals in polarity formation is far less understood. Biosensors are ideal tools for visualizing the regulation and function of biochemical pathways in organoids^64^, but their application has thus far achieved limited success^65^. The responses from kinase translocation reporters, which traffic from the nucleus to the cytosol upon phosphorylation by specific kinases, are difficult to quantify in such a compact volume^65^, whereas the modest dynamic ranges of FRET-based biosensors such as CKAR2 limit their sensitivity in 3D culture systems. With its dramatically enhanced dynamic range, ExRai-CKAR2 thus represents a felicitous development in our efforts to provide a continuous and quantifiable readout of endogenous aPKC activity in 3D culture. Using ExRai-CKAR2, we were able to detect basal aPKC activity in polarized and lumenized organoids, consistent with previous findings^22,51^. aPKCs are recruited to functionally distinct protein assemblies to drive cell polarity, but their exact spatiotemporal regulation is unclear^24,66^. By targeting ExRai-CKAR directly to Par6, we detected aPKC activity specifically at the Par complex and found that aPKC activity is enriched at the Par-complex, which is also shielded from cPKC and nPKC activities, an illustration of signaling compartmentation. Our new tools could benefit research in aPKC regulation as it cycles between functional complexes in more complex systems, such as patient-derived organoid models, using advanced organoid imaging technologies, including lattice lightsheet microscopy.

## Supporting information

ExRai-CKAR2 Supplementary

## Data availability

All data supporting the findings of this study are available upon reasonable request. Source data are provided with this paper.

## Code availability

Custom ImageJ macros and MATLAB code used to analyze imaging data are available on GitHub ^33^.

## Acknowledgments

The authors thank all members of the Zhang lab, especially Dr. Yanghao Zhong, for various technical help and discussion of the manuscript; Drs. Eric Griffis and Peng Guo of the UCSD Nikon Imaging Center for assistance with confocal imaging; Thomas Hoang for his technical help with cloning; Gema Lorden Losada from the A.C.N. lab for providing purified PKCα and technical support with *in vitro* characterization of ExRai-CKAR2. This work is supported by NIH grants R35 CA197622, R01 CA262815, and RF1 MH126707 (to J.Z.), R01 CA236386, R01 CA174869, and R01 CA262794 (to J.Y.), as well as by a TRDRP Postdoctoral Fellowship (T32FT5342) to Q.S.

## Author contributions

Q.S., S.M., J.Y., and J.Z. conceived the project. S.M. and J-F.Z. performed linker screening and generated ExRai-CKAR2. Q.S. performed *in vitro* characterization of ExRai-CKAR2, generated other constructs, and performed all live-cell and organoid imaging. Jing Zhang and Q.S. generated ExRai-CKAR2 and Cyto-ExRai-CKAR2 stable MDCK cell lines. Jing Zhang generated MDCK organoids. Q.S. generated HEK293T organoids. A.C.N., J.Y., and J.Z. supervised the project and coordinated experiments. Q.S. and Jing Zhang analyzed the data. Q.S., Jing Zhang, W.L., S.M., and J.Z. wrote the manuscript. Every author read and agreed on the final manuscript.

## Materials

Phorbol 12-myristate 13-acetate (PMA, LC Labs, P-1680), Phorbol 12,13-dibutyrate (PDBu, LC Labs, P-4833), Gö6983 (MilliporeSigma, G1918), Go6976 (MilliporeSigma, 365250), BAPTA-AM (Life Technologies, B6769), Thapsigargin (Cayman Chemical, 10522), Ionomycin (MilliporeSigma, 407951), B106 (Axon, # 2981, 98%), and pz09 (Reagency, #RNCY0048) were reconstituted in DMSO (Sigma-Aldrich) as 1000X stock solutions. UTP (Alfa, 98%+, J63427) was reconstited in ultrapure water (MilliQ). Lipofectamine 2000 (11668019) was purchased from Thermo Fisher. LysoTracker Red (L7528) was purchased from Thermo Fisher and diluted in DMSO.

## Plasmids

All primers used for molecular cloning are listed in Supplementary Table 1. To generate ExRai-CKAR linker variants, DNA fragments encoding cpEGFP, FHA1 and linkers were digested from pRRET-B ExRai-AKAR linker variants^31^ with SacI/EcoRI (Thermo Fisher, FD1134 and FD0275, respectively) and then ligated into a SacI/EcoRI-digested pRRET-B ExRai-CKAR1 backbone. pRSET-B ExRai-CKAR linker variants were then subcloned into pcDNA3 via BamHI/(Thermo Fisher FD1464) EcoRI digestion. ExRai-CKAR2 T/A was generated via Gibson Assembly using NEBuilder HiFi DNA Assembly Kit (New England Biolabs E2621) and primers 1–2. PM-, ER-, and lysosome-targeted ExRai-CKAR2 were made by subcloning a BamHI/EcoRI-digested fragment endocing full-length ExRai-CKAR2 into BamHI/EcoRI-digested pcDNA3 backbones containing the N-terminal 11 amino acids from Lyn (MGCIKSKRKDK), the N-terminal 27 amino acids from CYP450 (MDPVVVLGLCLSCLLLLSLWKQSYGGG), or full-length lysosome-associated membrane protein 1 (LAMP1). Cyto-ExRai-CKAR2 was constructed by subcloning a BamHI/XhoI-digested fragment into BamHI/XhoI-digested pcDNA3 containing a C-terminal nuclear export signal (LPPLERLTL). Par6-8aa-ExRai-CKAR2 was generated via Gibson Assembly of a Par6 fragment, PCR-amplified from pKMyc-Par6C (a gift from Ian Macara, Addgene plasmid # 15474; http://n2t.net/addgene:15474; RRID:Addgene_15474)^67^ using primers 3-4 to contain a BspEI and BamHI restriction sites and a GGTAGTGCTGGT spacer, into HindIII/BamHI-digested pcDNA3.1(+)-ExRai-CKAR2. A truncated EV49 linker containing BspEI/BamHI flanking sequences was PCR-amplified from EKAR4^42^ using primers 5-6 and inserted into BspEI/BamHI-digested Par6-ExRai-CKAR2 to obtain Par6-ExRai-CKAR2 with a 49-aa flexible linker. Lyso-DGK-mRuby2 was made by Gibson Assembly of DNA fragments encoding DGKζ (a gift from Stephen Gee, Addgene plasmid # 85454; http://n2t.net/addgene:85454; RRID:Addgene_85454)^68^ and mRuby2^69^, PCR-amplified using primers 7-10, into a BamHI/XbaI digested pcDNA3 backbone containing full-length LAMP1.

To make Lyso-Digda, the CFP of Digda was first replaced with ddRFP^35^ to reduce FP homology and facilitate further cloning. The DNA encoding ddRFP was amplified by PCR using primers 11-12, digested with EcoRI/XhoI, and ligated into Digda plasmid^44^ digested with the same enzymes. Then, the DNA fragment containing ddRFP-PKCb2C1-Venus-KRasCT and all ER/K rigid linkers was PCR-amplified using primers 13-14 to introduce a 3’ EcoRI site. A modified pcDNA3 backbone was PCR-amplified using primers 15-16 to remove the existing EcoRI site add a new EcoRI restriction site after XbaI. These two fragments were combined using Gibson Assembly to yield a modified ddRFP-PKCb2C1-Venus-KrasCT plasmid. LAMP1 was PCR-amplified using primers 17-18, followed by XbaI/EcoRI digestion and insertion into XbaI/EcoRI-digested ddRFP-PKCb2C1-Venus-KRasCT plasmid to replace the KRasCT. Cerulean (together with any 5’ extension exists in both parental and target plasmids) was PCR-amplified from Cer-FHA1-NES^70^ using primers 19-20, digested with HindIII/XhoI, and ligated into HindIII/XhoI-digested PKCb2C1-Venus-LAMP1.

For generation of MDCK stable cell lines, Cyto-ExRai-CKAR2 and Cyto-ExRai-CKAR2 T/A were PCR-amplified using forward primer aagcttgcggccgccacca and reverse primer gggccctctagattacag, digested with XbaI, and then ligated into EcoRV/XbaI-digested pLV-puro lentivirus vector. All plasmids were confirmed by DNA sequencing (Genewiz).

## Expression and purification ExRai-CKAR2

His-tagged ExRai-CKAR2 in pRSET-B was transformed into *E. coli* BL21 (DE3) chemically competent cells, and a single colony was inoculated in 20 mL LB-ampicilin (100 μg/ml) media and cultured overnight. The resulting seed culture was then used to inoculate 1 L of LB-ampicilin media. The culture was shaken vigorously (220 rpm) at 37 °C until reaching an OD_600_ of 0.6. ExRai-CKAR2 expression was induced by IPTG addition to a final concentration of 200 μM, followed by shaking at 18 °C for 20 h. Cells were harvested by centrifugation (5000 × g, 7 min, 4 °C) and stored at −80 °C until harvesting.

Frozen cell pellets were thawed on ice prior to resuspension in lysis buffer (50 mM Tris, pH 7.4, 300 mM NaCl, 10% glycerol, 20 mM imidazole) containing 1 mM PMSF and Complete EDTA-free Protease Inhibitor Cocktail (Roche). Cells were then lysed by probe sonication at 70% power and ten 30 s ON/30 s OFF cycles on ice. Cell debris was removed by centrifugation (45000 × g for 0.5 h, 4 °C), and the supernatant was loaded onto a pre-packed column with 5 mL Hispur^TM^ nickel nitrilotriacetic acid resin (Ni-NTA, GE Healthcare) through a syringe at roughly 1 ml/min. The column was washed with lysis buffer (20 mL), and washing continued with 50 ml washing buffer (50 mM Tris, pH 7.4, 300 mM NaCl, 10% glycerol, 60 mM imidazole) using syringes. The desired protein was eluted with lysis buffer supplemented to a total of 250 mM imidazole. Fractions were analyzed by SDS-PAGE, pooled, and concentrated using Amicon Ultra-15 centrifugal columns (30-kD cut-off, Millipore) and washed 3 times to exchange the buffer to 50 mM Tris, pH 7.4, 300 mM NaCl, 10% glycerol. Protein concentrations were determined using the Pierce BCA Protein Assay Kit (Thermo Fisher Scientific) on a Spark 20M microplate reader (Tecan).

*In vitro* excitation and emission spectra were obtained on a PTI QM-400 fluorometer using FelixGX v.4.1.2 software (Horiba). Excitation scans were collected at 530 nm emission, and emission scans were performed at 400- and 480 nm excitation. Purified biosensor (1 μM) was incubated with 5 μg of PKCα for 30 min at 30 °C in kinase assay buffer (20 mM HEPES, pH 7.4, 2 mM DTT, 5 mM MgCl2, 140 µM/3.8 µM phosphatidylserine/diacylglycerol membranes) without ATP or lipids and with 500 µM EGTA (no Ca^2+^) for unphosphorylated spectra, or with 100 µM ATP and 100 µM Ca^2+^ (no EGTA) for phosphorylated spectra.

## Cell culture and transfection

HeLa and Cos7 cells were cultured in Dulbecco’s modified Eagle medium (DMEM) (Gibco) containing 1 g l^−1^ or 4.5 g l^−1^ glucose respectively, 10% (v/v) fetal bovine serum (FBS) (Sigma), and 1% (v/v) penicillin-streptomycin (Pen-Strep) (Sigma-Aldrich). MDCK cells were cultured in Minimum Essential Medium Eagle (MEM) (Sigma) containing 10% (v/v) fetal bovine serum (FBS) (Sigma) and 1% (v/v) penicillin-streptomycin (Pen-Strep) (Sigma-Aldrich).

All cells were maintained at 37 °C in a humidified atmosphere of 5% CO_2_. For imaging experiments, cells were plated onto sterile 35-mm glass-bottomed dishes and grown to 50–70% confluence. Transient transfection was performed using Lipofectamine 2000 (Invitrogen), after which cells were cultured for an additional 24 h (HeLa, Cos7).

## Biosensor localization

Cos7 cells expressing Lysosome- or ER-targeted ExRai-CKAR2 were stained for 30 min with LysoTracker RED (Invitrogen) at a final concentration of 1 mM in Hank’s balanced salt solution (HBSS) (Gibco) or co-expressed with ER-mRuby2, respectively. Cells were imaged on a Nikon Ti2 spinning-disk confocal microscope (Nikon Instruments) equipped with an SR HP APO TIRF 100x/1.49 numerical aperture (NA) oil objective (Nikon), excitation disk CSU-X1 (Yokogawa) under SoRa mode, and a Photometrics Prime95B sCMOS camera (Photometrics) controlled by NIS-Elements software (High content analysis package, Nikon). ExRai-CKAR2 images were acquired using 488- and 405-nm lasers at 15 and 20% power, respectively, and 200-ms exposure times. Red fluorescent protein (RFP) images were acquired using a 561-nm laser at 20% power and 200-ms exposure time.

## MDCK stable cell line generation and organoid culture

MDCK stable cell lines were generated using lentiviral plasmid vectors. Briefly, 293T cells were transfected with pCMVΔ8.2R, VSVG and pLV-Puro lentiviral construct expressing Cyto-ExRai-CKAR2 or Cyto-ExRai-CKAR2 T/A. Viral supernatants were then concentrated using Lenti-X^TM^ concentrators (Takara). Concentrated viral supernatants were applied to MDCK cells with 6 µg/ml protamine sulfate. Infection was repeated the next day. Infected cells were then selected with puromycin (2 µg/ml).

MDCK cells were grown in 3D culture as previously described^71^. Briefly, Matrigel (Growth Factor Reduced Matrigel, BD Biosciences) was plated onto 35-mm glass-bottom dish and allowed to solidify for 30 min at 37 °C. MDCK cells were trypsinized, resuspended in media supplemented with 2% Matrigel and then plated on top of Matrigel.

## Time-lapse fluorescence imaging

Cells were washed twice with HBSS supplemented with 20 mM HEPES, pH 7.4, and 2 g/l and subsequently imaged in the same buffer in the dark at 37 °C. PMA, PDBu, UTP, Gö6983, Go6976, or B106 were added as indicated.

HeLa and Cos7 cells were imaged on a Zeiss AxioObserver Z7 microscope (Carl Zeiss) equipped with a Definite Focus.2 system (Carl Zeiss), a 40×/1.4 NA oil objective and a Photometrics Prime95B sCMOS camera (Photometrics) controlled by METAFLUOR 7.7 software (Molecular Devices). Dual GFP excitation-ratio imaging was performed using 480DF20 and 405DF20 excitation filters, a 505DRLP dichroic mirror and a 535DF50 emission filter. RFP intensity was imaged using a 572DF35 excitation filter, a 594DRLP dichroic mirror and a 645DF75 emission filter. Dual cyan/yellow emission ratio imaging was performed using a 420DF20 excitation filter, a 455DRLP dichroic mirror and two emission filters (473DF24 for cyan fluorescent protein and 535DF25 for yellow fluorescent protein). Filter sets were alternated by an LEP MAC6000 control module (Ludl Electronic Products Ltd). Exposure times ranged from 100 to 500 ms, and images were acquired every 30 s. MDCK cells were imaged on the same system, controlled by a modified version of the open-source MATLAB (Mathworks) and μmanager (Micro-Manager)-based MATScope imaging suite ^33^.

MDCK organoids were imaged on a Nikon Ti2 spinning-disk confocal microscope (Nikon Instruments) equipped with a CFI Apo LWD Lambda S 40XC/1.15 numerical aperture (NA) water immersion objective (Nikon), excitation disk CSU-X1 (Yokogawa) under SoRa mode, and a Photometrics Prime95B sCMOS camera (Photometrics) controlled by NIS-Elements software (High content analysis package, Nikon). ExRai-CKAR2 images were acquired using 488- and 405-nm lasers at 20 and 15% power, respectively, with 500-ms exposure times every 45s. Raw fluorescence images were corrected by subtracting the background fluorescence intensity of a cell-free region from the emission intensities of biosensor-expressing cells at each time point. For comparison among CKAR2, ExRai-CKAR1 and ExRai-CKAR2, cytosolic regions-of-interest (ROIs) are selected for analysis. GFP excitation ratios (Ex480/Ex405) and RFP fluorescence intensities were then calculated at each time point. All biosensor response time courses were subsequently plotted as the normalized fluorescence intensity or ratio change with respect to time zero (ΔF/F_0_ or ΔR/R_0_), calculated as (F − F_0_)/F_0_ or (R − R_0_)/R_0_, where F and R are the fluorescence intensity and ratio value at a given time point, and F_0_ and R_0_ are the initial fluorescence intensity or ratio value at time zero, which was defined as the time point immediately preceding drug addition. Maximum intensity (ΔF/F) or ratio (ΔR/R) changes were calculated as (F_max_ − F_min_)/F_min_ or (R_max_ − R_min_)/R_min_, where F_max_ and F_min_ or R_max_ and R_min_ are the maximum and minimum intensity or ratio value recorded after stimulation, respectively. Graphs were plotted using GraphPad Prism 8 (GraphPad Software).

## Statistics and reproducibility

All experiments were independently repeated as noted in the figure legends. Statistical analyses were performed using GraphPad Prism 8. For Gaussian data, Student’s t test was used for pairwise comparisons. For comparing three or more sets of data, ordinary one-way ANOVA followed by multiple comparisons test was performed. Nonparametric tests were performed using U-test for pairwise comparisons or the Kruskal– Wallis test followed by Dunn’s multiple comparisons test for analyses of three or more groups. Statistical significance was defined *as P* < 0.05 with a 95% confidence interval. The number of cells analyzed (n cells) and number of independent experiments are reported in all figure legends. All time courses and dot plots shown depict the mean ± standard error of the mean, and violin plots depict the median and quartiles, unless otherwise noted.

