## Supplementary material for "Sensitive Fluorescent Biosensor Reveals Differential Subcellular Regulation of PKC": ExRai-CKAR2 Supplementary

**Supplementary Figure 1**
**Supplementary Figure 2**
**Supplementary Figure 3**
**Supplementary Figure 4**
**Supplementary Figure 5**
**Supplementary Figure 6**
**Supplementary Figure 7**
**Supplementary Figure 8**
**Supplementary Figure 9**
**Supplementary Figure 10**
**Supplementary Figure 11**
**Supplementary Figure 12**
**Supplementary Figure 13**
**Supplementary Table 1**

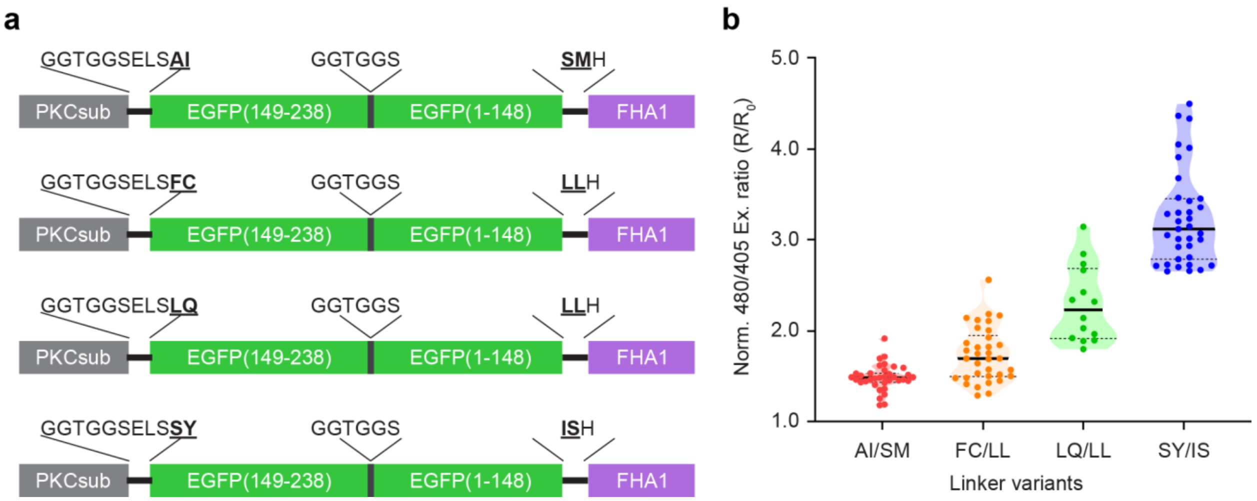


**Supplementary Figure 1 | ExRai-CKAR linker optimization**. (Left) Domain structure of each linker variant. (Right) Quantification of the maximum excitation ratio (480/405) responses to stimulation with 50 ng/ml PMA in HeLa cells expressing ExRai-CKAR linker variants .

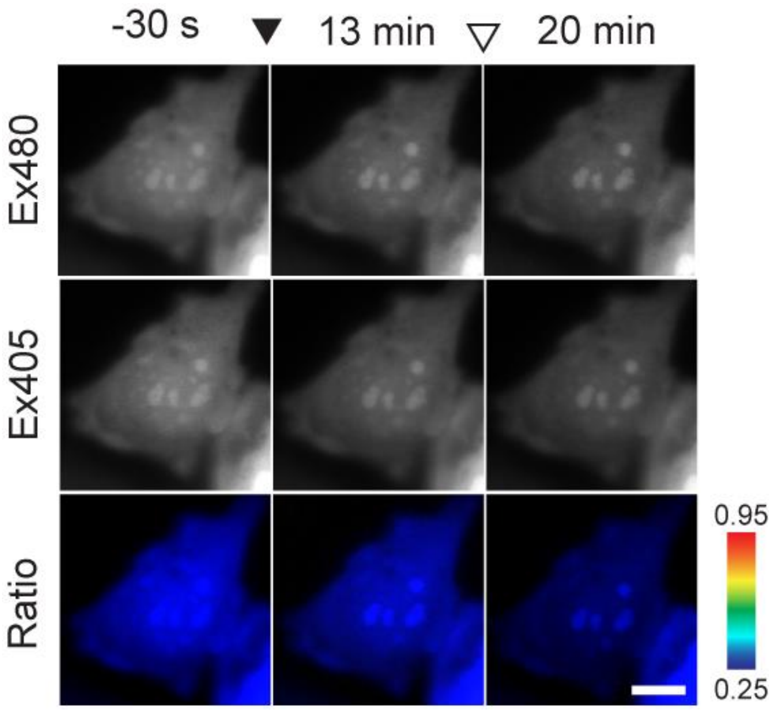


**Supplementary Figure 2 |** Representative images showing ExRai-CKAR2 T/A fluorescence at 480-nm (Ex480, top) and 405-nm (Ex405, middle) excitation and pseudocolored images (bottom) showing the excitation ratio response to PMA and Gö6983 addition in Cos7 cells. The solid arrowhead indicates time PMA addition, and the hollow arrowhead indicates Gö6983 addition. Images are representative of three independent experiments. Warmer colors indicate higher ratios. Scale bar, 10 μm.

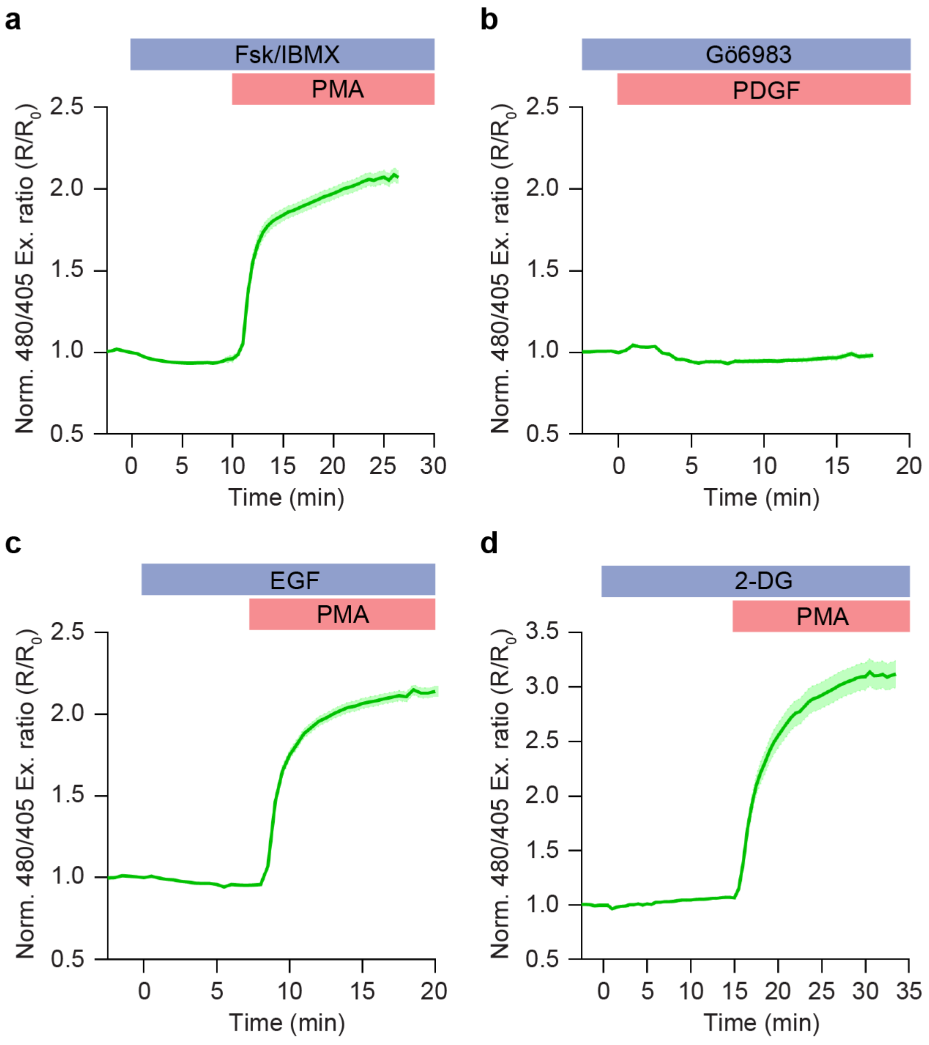


**Supplementary Figure 3 | Specificity of ExRai-CKAR2.** Average time courses excitation ratio (480/405) response from ExRai-CKAR2 in (a) HeLa cells stimulated with Forskolin and 3-isobutyl-1-methylxanthine (Fsk-IBMX) followed by PMA; (b)NIH3T3 cells pretreated for 30 min with Gö6983 followed by platelet-derived growth factor (PDGF) stimulation; (c) HEK293T cells stimulated with epidermal growth factor (EGF) followed by PMA; and (d) Cos7 cells stimulated with 2-deoxy glucose (2-DG) followed by PMA. Prior to imaging, cells in **b** were serum-starved fro 18 h, and cells in **c** and **d** were pre-incubated in HBSS imaging buffer for 30 min at 37 °C. Time courses are representative of three independent experiments; solid lines indicate mean responses; shaded areas, s.e.m.

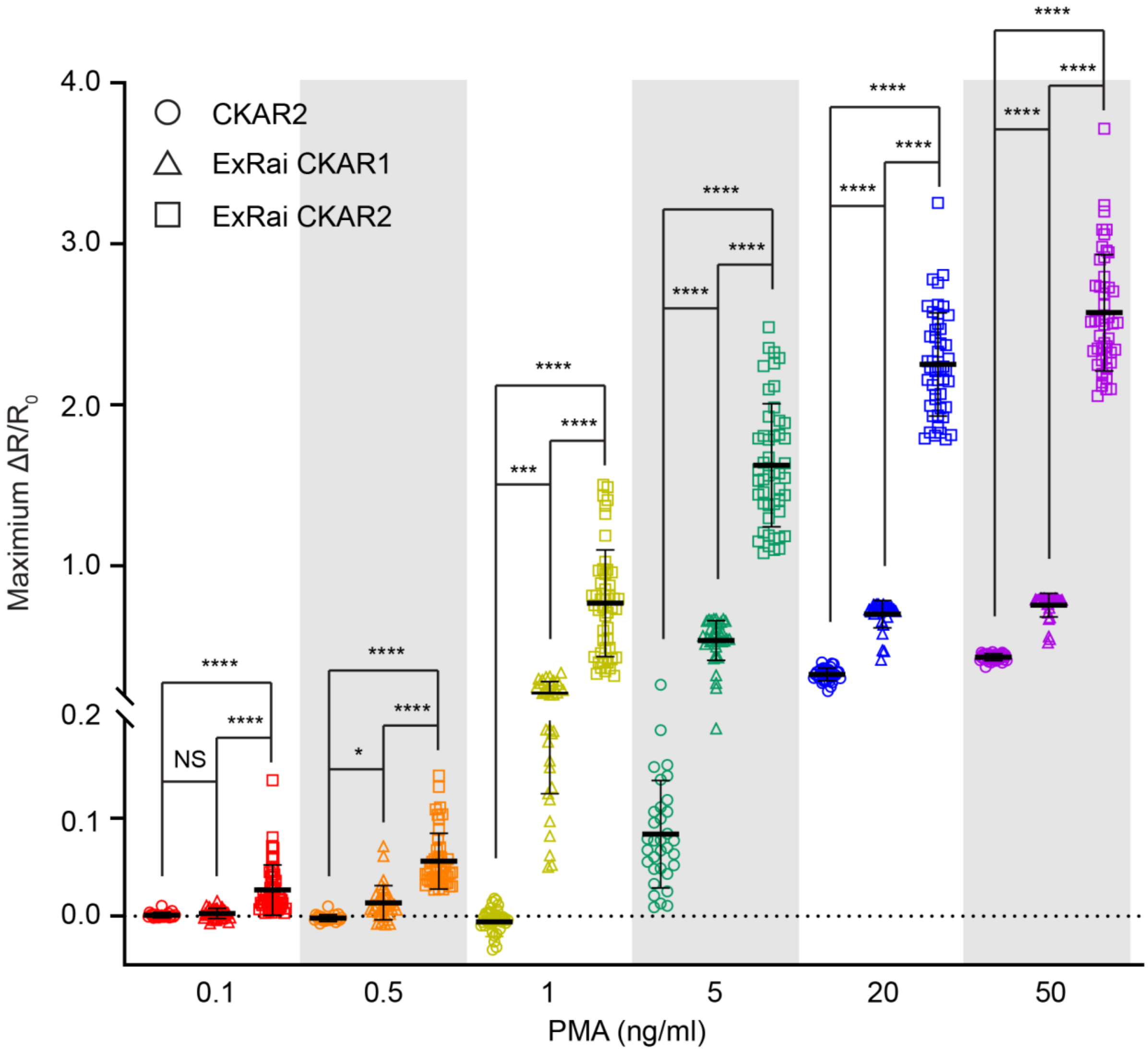


**Supplementary Figure 4 | Dose-dependent response of PKC biosensors.** Quantification of maximum yellow/cyan emisison ratio or 480/405 excitation ratio responses ($\Delta R/R_0$) of CKAR2, ExRai-CKAR1, and ExRai-CKAR2 in HeLa cells stimulated with increasing concentrations of PMA. For 0.5 ng/ml PMA treatment, **P* = 0.0271; For 1 ng/ml PMA treatment, ****P* = 0.0001. Statistical analysis was performed using ordinary one-way ANOVA followed by Dunnett's multiple-comparison test. *****P* < 0.0001, NS, not significant. Data are mean ± s.e.m.

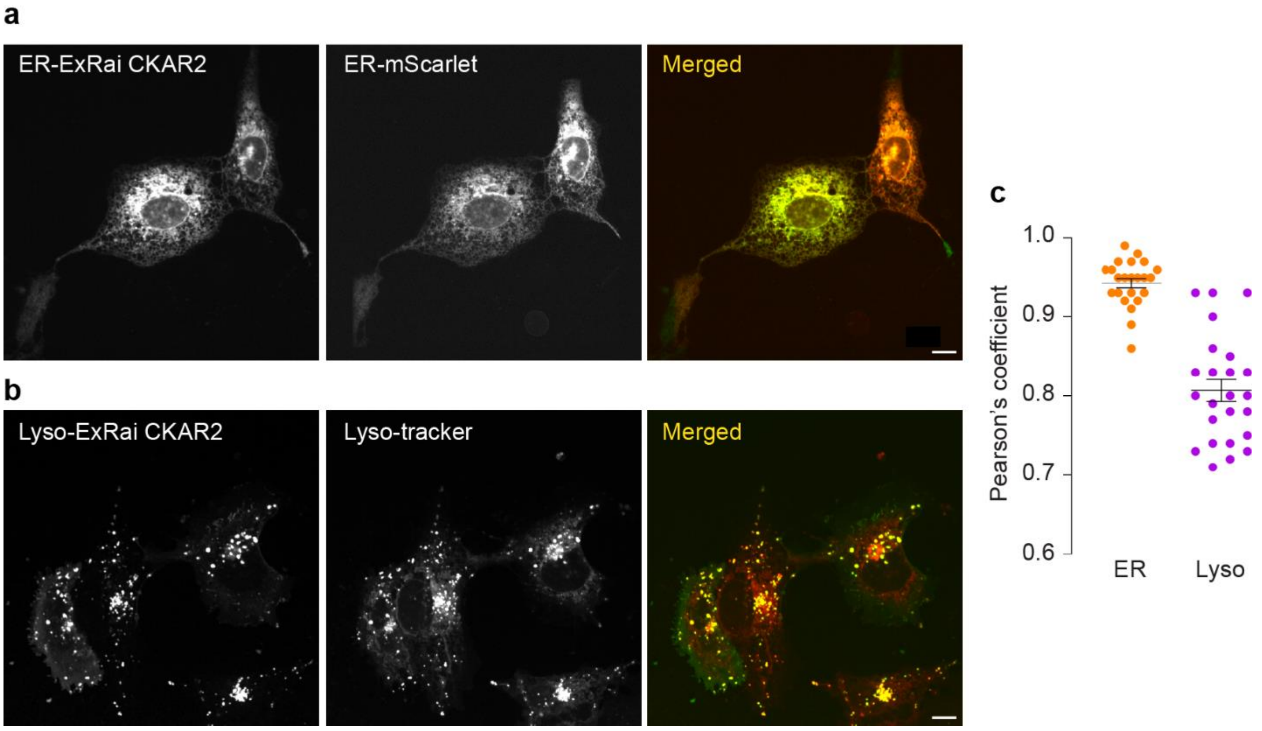


**Supplementary Figure 5 | Colocalization of ER-ExRai-CKAR2 and Lyso-ExRai-CKAR2 with corresponding organelle markers. a**, Representative images of ER-ExRai-CKAR2 (Ex480 channel), ER-mScarlet, and merged image. **b**, Representative images of Lyso-ExRai-CKAR2 (Ex480 channel), LysoTracker-Red, and merged image. Scale bars, 10 µm. **c**, Quantification of Pearson's coefficient of ER-ExRai-CKAR2 and ER-mScarlet (n = 23 cells) and Lyso-ExRai-CKAR2 and LysoTracker-Red (n = 24 cells). Data are mean ± s.e.m.

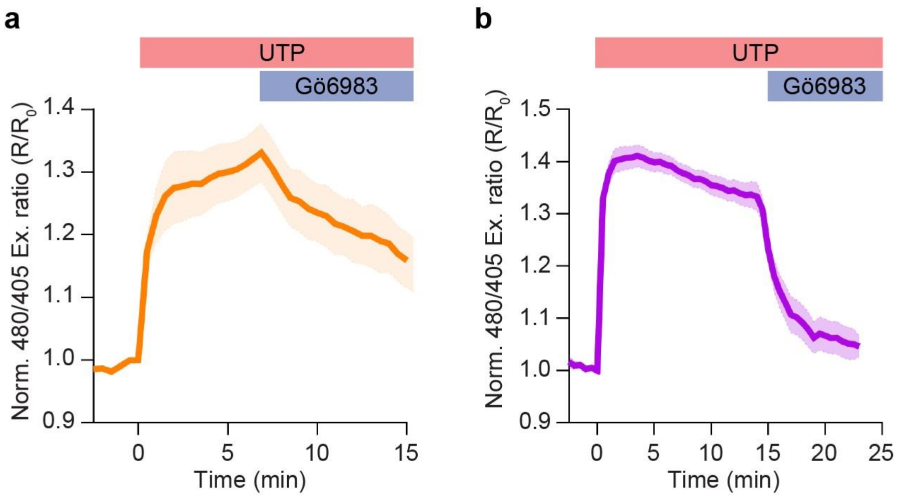


**Supplementary Figure 6 |** Average time courses of normalized excitation ratio (480/405) responses from (a) ER-ExRai-CKAR2 and (b) Lyso-ExRai-CKAR2 in Cos7 cells treated with 100 μM UTP followed by the pan-PKC inhibitor Gö6983. Time courses are representative of three independent experiments; solid lines indicate mean responses; shaded areas, s.e.m.

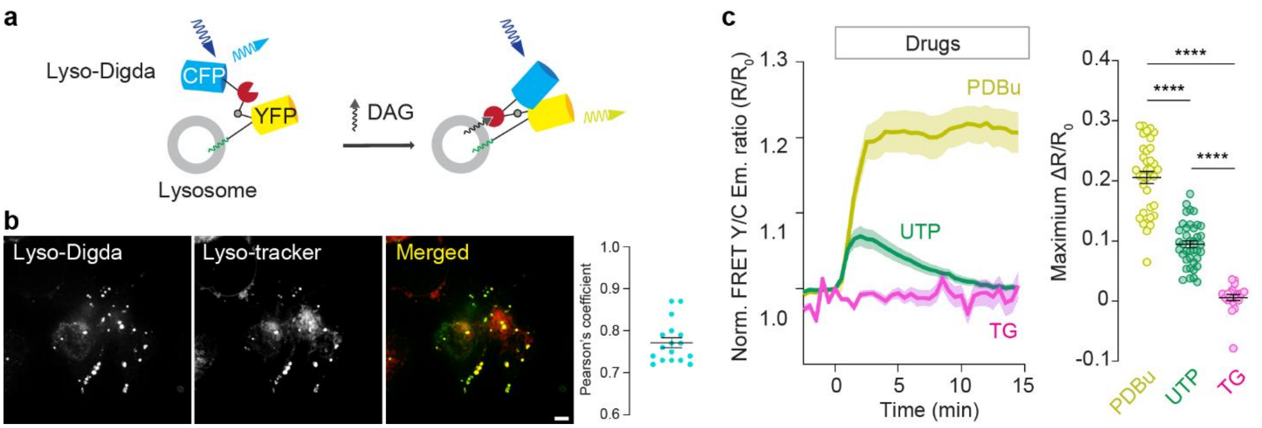


**Supplementary Figure 7 | Development and characterization of Lyso-Digda. a**, Schematic of lysosome-targeted DAG biosensor Lyso-Digda. Lyso-Digda contains the DAG-binding C1 domain of PKCβII sandwiched between cyan (Cerulean) and yellow (mVenus) fluorescent proteins through rigid α-helical linkers consisting of repeated EAAAR sequences. A di-glycine motif serves as a flexible hinge in the middle. The biosensor is targeted to the lysosomal membrane by tethering full-length LAMP1to the rigid α-helical linker at the C-terminus. Upon DAG stimulation, the C1 domain binds to DAG generated on the lysosomal membrane, inducing a conformational change in the biosensor through the flexible di-glycine motif that brings the CFP into proximity with YFP. **b**, Colocalization of Lyso-Digda with LysoTracker-Red. (left) Representative images of Lyso-Digda, LysoTracker-Red, and merged image. (right) Quantification of the Pearson's coefficient from n = 17 cells. Data are mean ± s.e.m. **c**, Average time courses (left) of normalized yellow/cyan emission ratio responses from Lyso-Digda in Cos7 cells stimulated with 200 nM PDBu (n = 35 cells), 100 μM UTP (n= 35 cells) or 1 μM TG (n= 21 cells). Quantification of maximum responses (right) from n = 35, n = 35 and n = 21 cells from three independent experiments. All time courses are representative of three independent experiments; solid lines indicate mean responses; shaded areas, s.e.m. Statistical analysis was performed using ordinary one-way ANOVA followed by Tukey's multiple-comparison test. ****$P < 0.0001$. Data are mean ± s.e.m.

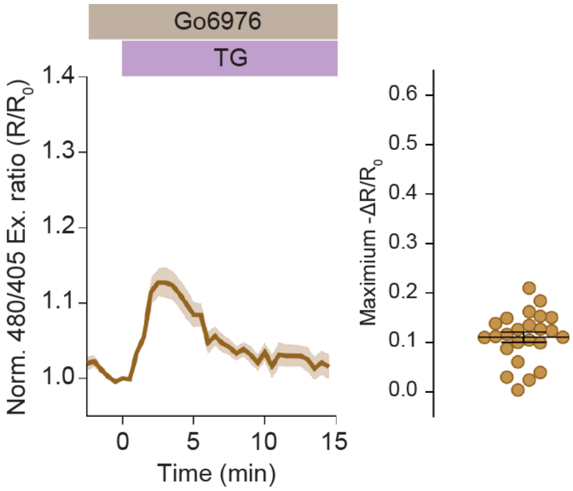


**Supplementary Figure 8 |** Representative average time courses (left) of normalized excitation ratio (Ex480/405) response from Lyso-ExRai-CKAR2 in Cos7 cells pretreated for 30 min with the cPKC inhibitor Go6976 (n = 11 cells) followed by stimulation with thapsigargin (TG). Quantification of maximum response (right), n = 24 cells from three independent experiments.

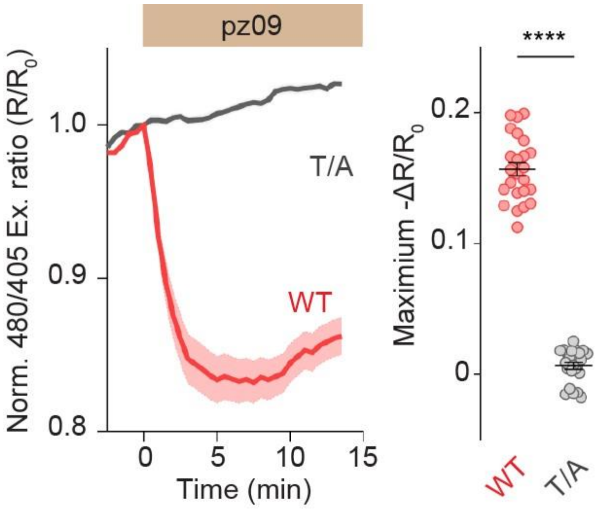


**Supplementary Figure 9 | Response of Cyto-ExRai-CKAR2 in 2D-cultured MDCK cells.** Representative average time courses (left) of normalized excitation ratio (Ex480/405) responses from Cyto-ExRai-CKAR2 (WT, n = 24 cells) and the negative control Cyto-ExRai CKAR2 T/A (T/A, n = 23 cells) in MDCK cells treated with pz09. Quantification of maximum response (right) from n = 24 and n = 23 cells from three independent experiments. Time courses are representative of three independent experiments; solid lines indicate mean responses; shaded areas, s.e.m. ****P < 0.0001 according to Student's *t*-test. Data are mean ± s.e.m.

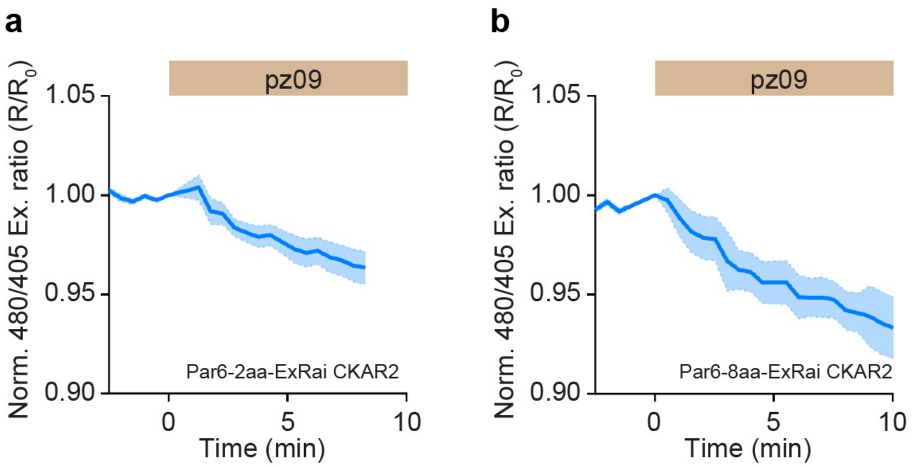


**Supplementary Figure 10 |** Average time courses of normalized excitation ratio (480/405) respones from Par6-2aa-ExRai-CKAR2 and Par6-8aa-ExRai-CKAR2 in HeLa cells treated with pz09. n = 17 cells each variant from 3 (2 aa) and 4 (8aa) independent experiments. Solid lines indicate mean responses; shaded areas, s.e.m.

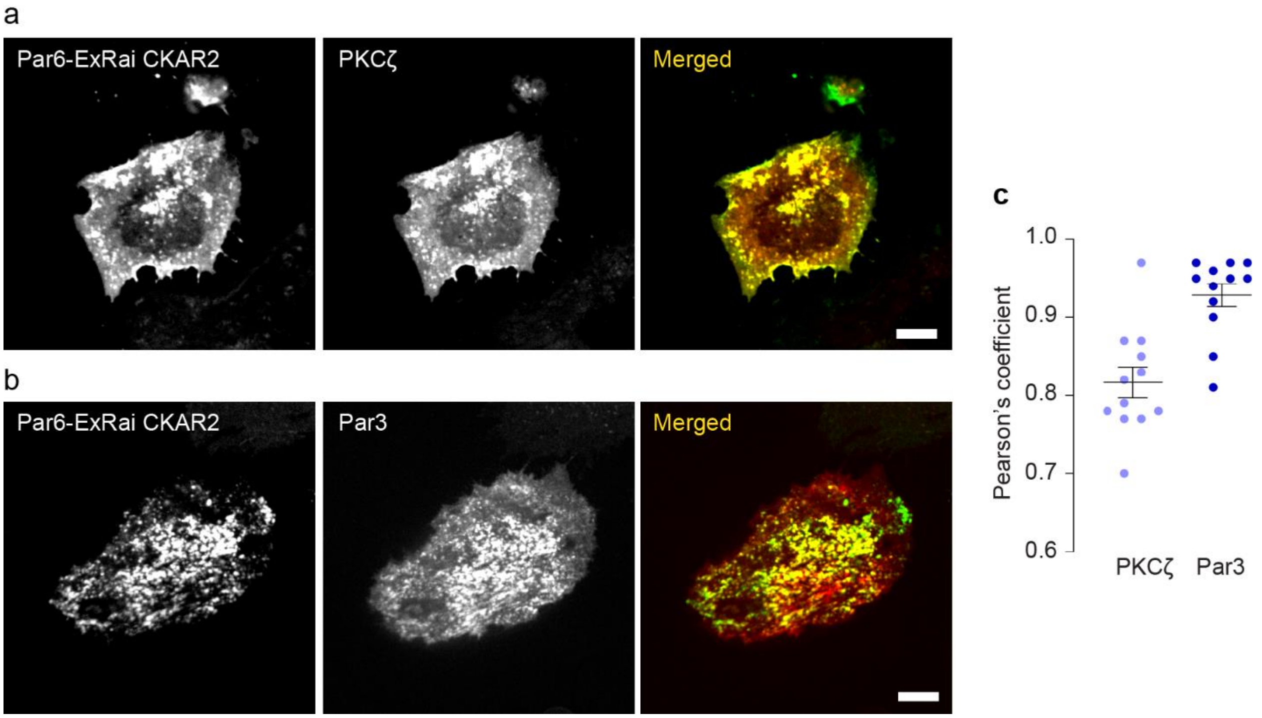


**Supplementary Figure 11 | Colocalization of Par6-ExRai CKAR2 with Par3 and PKCζ. a**, Representative images of HeLa cells co-expressing Par6-ExRai-CKAR2 and PKCζ -mCherry. **b**, Representative images of HeLa cells co-expressing Par6-ExRai-CKAR2 and Par3-mCherry. Scale bars, 10 μm. **c**, Quantification of the Pearson's coefficient of Par6-ExRai-CKAR2 and PKCζ -mCherry (n = 12 cells) and Par3-mCherry (n = 12 cells). Data are mean ± s.e.m.

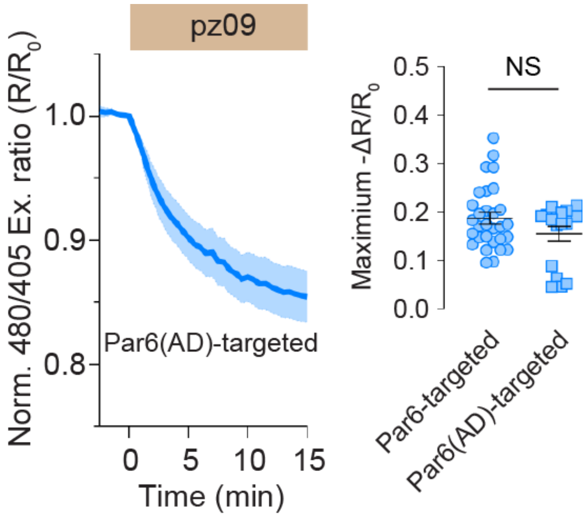


**Supplementary Figure 12 | Response of Par6(AD)-ExRai-CKAR2 in HEK293T cells.** Representative average time course (left) of normalized excitation ratio (Ex480/405) responses from Par6(AD)-ExRai-CKAR2 ($n = 9$ cells) in HEK293T cells treated with pz09. Quantification of maximum response (right) from $n = 30$ and $n = 18$ cells from three independent experiments. Time course is representative of three independent experiments; solid lines indicate mean responses; shaded areas, s.e.m. NS, not significant according to Student's *t*-test. Data are mean ± s.e.m.

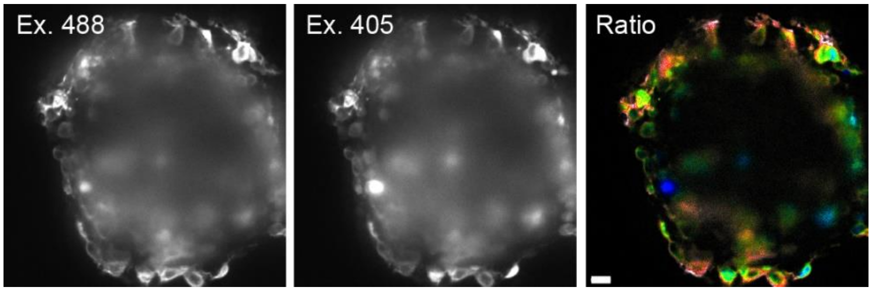


**Supplementary figure 13 |** Fluorescence and pseudocolor images of a representative 3D-cultured HEK293T organoid expressing Par6-ExRai-CKAR2. Scale bar, 20 µm.

Supplementary Table 1. Primers used in this study

| **Primer** | **Primer name** | **Primer sequence** |
|---|---|---|
| **1** | ExRaiCKAR2 T/A Foward | GCGATTCAGAAGATTCCAGGCGTTGAAGGACAAGGCTAAGG |
| **2** | ExRaiCKAR2 T/A Reverse | CCTTAGCCTTGTCCTTCAACGCCTGGAATCTTCTGAATCGC |
| **3** | Par6 Forward | tggctagcgtttaaacttAAGCTTATGGCCCGGCCGCAG |
| **4** | Par6 Reverse | TACCACCAGCACTGGTACCTCCGGAGAGGCTGAAGCCACTACCATCTCCTC |
| **5** | EV Forward | TGGTAGTGGCTTCAGCCTCTCCGGAGGTACCAGTGCTGGTGG |
| **6** | EV_Bam Reverse | GGAATCTTCTGAATCGCATggatccACCACCAGCACTACCAC |
| **7** | DGKζ Forward | tcggtggtggcggtggtgcggatccccgggacggtagcc |
| **8** | DGKζ Reverse | gacaccatgaattccacagccgtctcctg |
| **9** | mRuby2 Forward | gctgtggaattcatggtgtctaagggcg |
| **10** | mRuby2 Reverse | acactatagAATAGGGCCCTCTAGAttacttgtacagctcgtcca |
| **11** | ddRFP Forward | aaattgaattcggcATGGTGAGCAAGAGCGAG |
| **12** | ddRFP Reverse | ttaaactcgagGCTGCCGGTGCCATG |
| **13** | Bam-ddRFP Forward | ACGATGACGATAAggatcccATGGTGAGCAAGAGCGAG |
| **14** | KRasCT-EcoR_Reverse | AATAGGGCCCGaattCTTACATAATTACACACTTTGTCTTTGACTTC |
| **15** | pcDNA_Forward | GTAAGaattCGGGCCCTATTctatagtgtcacc |
| **16** | pcDNA_Reverse | gggatccTTATCGTCATCGTC |
| **17** | Lamp1_Forward | aaatttctagaACCATGGCGGCGC |
| **18** | Lamp1_Reverse | ttaaaGaattCTTAgatagtctggtagcctgcgtga |
| **19** | HindIII Forward | actcactatagggAGACCCAAG |
| **20** | Cer_Reverse | GCCTCTCTAGCTGCGGCTTCctcgagCTTGTACAGCTCGTCCATG |
